# State-Dependent Transcriptomic Collapse of the Brain’s Lactate and Ketone Thermodynamic Sensors in Schizophrenia

**DOI:** 10.64898/2026.06.26.734782

**Authors:** Bryan A. Krantz

## Abstract

Metabolic psychiatry has recently achieved unprecedented clinical rescue in treatment-resistant Schizophrenia (SCZ) utilizing targeted ketogenic interventions. However, the field has operated without a defined genomic anchor, leaving the biophysical mechanism of these therapies largely unexplained. Here, we report the discovery of the definitive metabolic sensor array driving this pathology. By integrating high-resolution topological mapping of SCZ GWAS summary statistics, 3D chromatin conformation (Hi-C), and multi-tissue transcriptomics, we identify massive, non-coding structural variances flanking the *HCAR2/HCAR1* tandem locus—the brain’s master thermodynamic governor. We demonstrate that while the protein-coding hardware of these receptors remains intact, their shared 3D Topologically Associating Domain (TAD) is fundamentally fractured. This structural collapse drives a perfect transcriptomic double dissociation in the human cortex: the 3’ mutational “skyscraper” severely downregulates the *HCAR1* lactate emergency brake, while the 5’ mutational cluster selectively paralyzes the *HCAR2* β-hydroxybutyrate (BHB) and niacin cooling switch. This dual-flank enhancer failure elegantly provides a definitive genomic etiology for historical SCZ biomarkers, physically explaining both chronic cerebrospinal fluid lactate pooling and the infamous “absent niacin flush.” Furthermore, peripheral eQTL mapping reveals profound antagonistic pleiotropy, characterized by a hyper-activation of the *HCAR1* lactate shuttle in the testis, explaining the evolutionary conservation of this metabolically catastrophic architecture. Ultimately, we reframe Schizophrenia not as an intrinsic neurological defect, but as an evolutionary “fuel mismatch.” The high-performance cognitive architecture of the hominid brain, evolved for ancestral ketogenic environments, experiences a catastrophic thermodynamic crash when deprived of its requisite BHB coolant by modern, high-glycemic diets.

## Introduction

Despite decades of intense focus on dopaminergic neurotransmission, the physiological hallmarks of Schizophrenia (SCZ) strongly indicate a macroscopic thermodynamic failure. Clinical and neuroimaging observations consistently report localized, chronic lactate pooling within the central nervous system ^1^, suppressed systemic glycolytic shifting, and an absent or profoundly blunted dermal “niacin flush” response ^2^. Furthermore, emerging metabolic interventions—such as ketogenic therapies that elevate systemic β-hydroxybutyrate (BHB)—have demonstrated unprecedented clinical rescue in treatment-resistant SCZ cohorts ^3,4^. Yet, the field lacks a unified biophysical model linking the population-level genetic risk of SCZ to these specific, system-wide failures in energy regulation, lactate shuttling, and metabolic cooling.

During periods of intense cognitive demand and extreme energy utilization, transient Pyruvate Dehydrogenase (PDH) glycolytic blockade can occur, leading to a rapid, localized accumulation of unburned lactate. Rather than merely a metabolic waste product, this lactate serves as a critical signaling molecule. The brain manages this intense energetic flux via the Hydroxycarboxylic Acid Receptor (HCAR) family—a uniquely conserved class of 7-transmembrane helix G-protein coupled receptors (GPCRs) that act as master sensors of cellular thermodynamic status ^5^. Within this network, *HCAR1* (the lactate receptor) acts as a localized emergency brake: as extracellular lactate pools during glycolytic panic, *HCAR1* activation suppresses cAMP production, hyperpolarizes the membrane, and throttles down neuronal firing to prevent excitotoxicity ^6^. Conversely, the adjacent *HCAR2* (the BHB and niacin receptor) functions as a systemic cooling switch, detecting ketogenic substrates to trigger anti-inflammatory ^7^, lipid-mobilizing cascades that restore baseline metabolic homeostasis.

Standard genome-wide association studies (GWAS) have repeatedly implicated broad metabolic and calcium-channel loci in SCZ ^8^, but bulk statistical pipelines often obscure the localized regulatory architecture driving these signals. By performing high-resolution topological mapping of Psychiatric Genomics Consortium (PGC) summary statistics, we isolated a critical, tandemly duplicated vulnerability on Chromosome 12 corresponding exactly to this *HCAR2* and *HCAR1* array, which functions collectively as the brain’s baseline thermodynamic governor. Strikingly, we observed that the coding sequences for both receptor bodies remain structurally intact; the SCZ mutational burden is driven entirely by massive structural variances localized strictly to the distal 3’ and 5’ non-coding regulatory flanks.

In this study, we bridge population genetics with tissue-specific transcriptomics to demonstrate that this non-coding architectural variance induces a perfect transcriptomic double dissociation. Utilizing targeted eQTL mapping across the human cortex and highly metabolic peripheral tissues, we reveal that the 3’ mutational “skyscraper” independently and severely downregulates the *HCAR1* lactate emergency brake, while the 5’ mutational cluster selectively paralyzes the *HCAR2* cooling switch. This dual-flank enhancer failure effectively shears off the brain’s metabolic governors, leaving the cortical network locked in a state of runaway glycolytic panic and providing a definitive genomic etiology for the thermodynamic hallmarks of Schizophrenia.

## Results

### The Lactate Signaling Network and Genomic Topology of the HCAR Locus

We hypothesized that aberrant lactate signaling within the cortex is a fundamental driver of SCZ pathology, representing a macroscopic thermodynamic failure rather than a strictly localized neurotransmitter imbalance. To investigate the genomic architecture of this metabolic crisis, we systematically probed Psychiatric Genomics Consortium (PGC) Wave 3 summary statistics ^9^ for structural variance across major factors governing lactate biochemistry and cellular signaling (**Table 1**). Specifically, we evaluated the regional mutational burdens of the core metabolic enzymes (*LDHA*, *LDHB*), the transcription factor driving activity-dependent rewiring (*CREB1*), the master regulator of the glycolytic shift (*HIF1A*), the metabolic proto-oncogene (*MYC*), and the neuronal lactate receptor (*HCAR1*). This network-wide analysis revealed significant regional genomic vulnerabilities distributed across these core thermodynamic and plasticity loci (**Figure 1A, Supplemental Figure S1)**.

**Figure 1.**
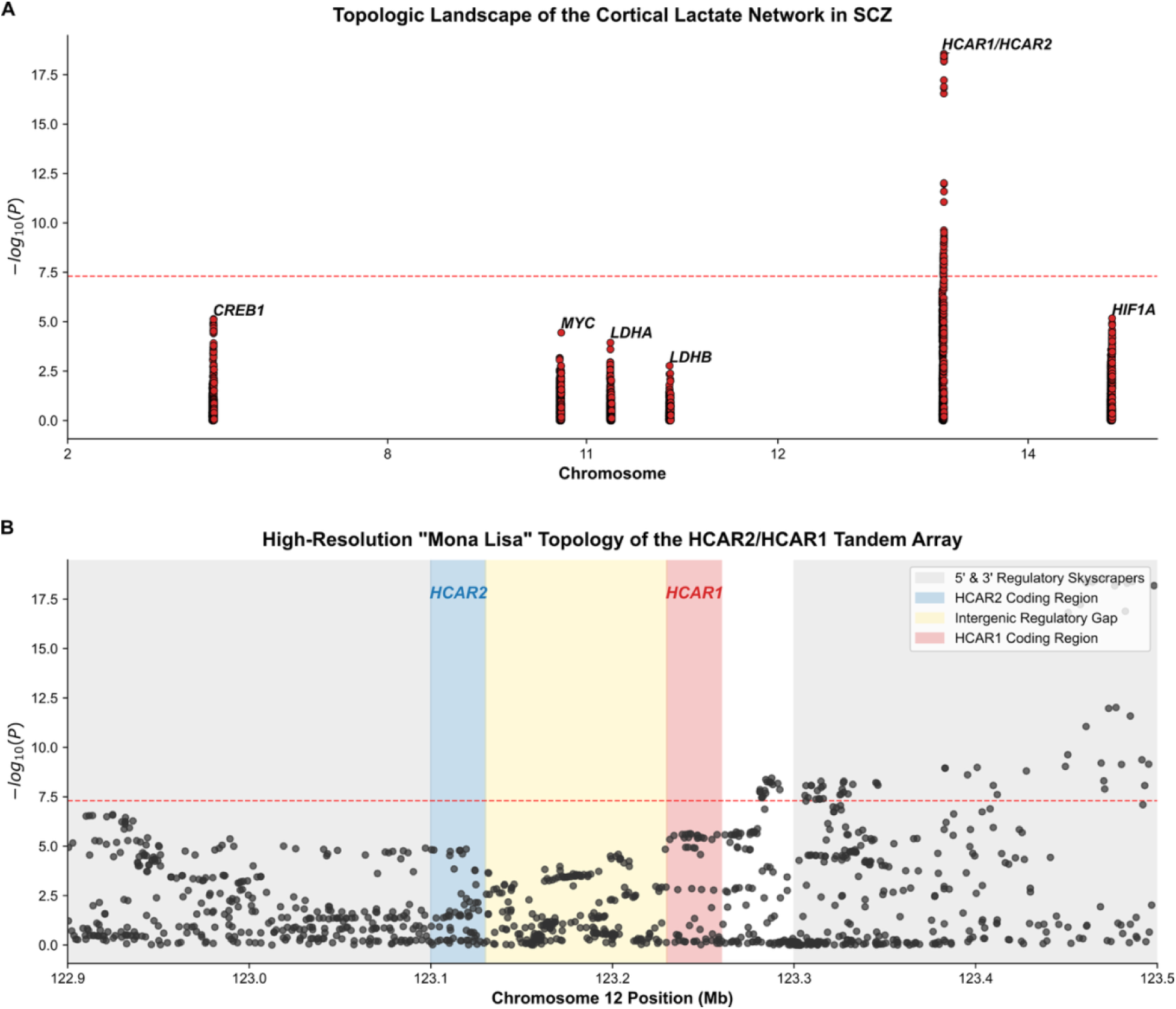
The Lactate Network and the HCAR Topological Locus Plot. **(A)** Topologic Landscape of the Cortical Lactate Network. A targeted Manhattan Plot highlighting the significant regional mutational burdens across major lactate biochemistry and signaling loci (*LDHA*, *LDHB*, *CREB1*, *HIF1A*, *MYC*, and the *HCAR1/HCAR2* tandem array) in the Schizophrenia (SCZ) cohort. The dashed red line denotes standard genome-wide significance (*P* = 5 × 10^-8^). **(B)** High-Resolution Topology of the *HCAR2/HCAR1* Tandem Array. A Locus-Zoom plot of the Chromosome 12 region (122.9 – 123.5 Mb) isolating the specific regulatory architecture of the tandem metabolic governor. The actual coding sequences for both *HCAR2* (blue shaded region; BHB cooling switch) and *HCAR1* (red shaded region; lactate emergency brake) harbor virtually zero significant structural variance (flat, non-significant *P*-values). Instead, they are bookended by towering mutational “skyscrapers” at the 3’ and 5’ regulatory extremities (grey shaded regions), separated by an intergenic gap (yellow). This structural asymmetry demonstrates that the physical hardware of the brain’s metabolic governors remains perfectly intact, but the shared 3D non-coding enhancer network dictating their joint expression is fundamentally fractured.

**Table 1.**
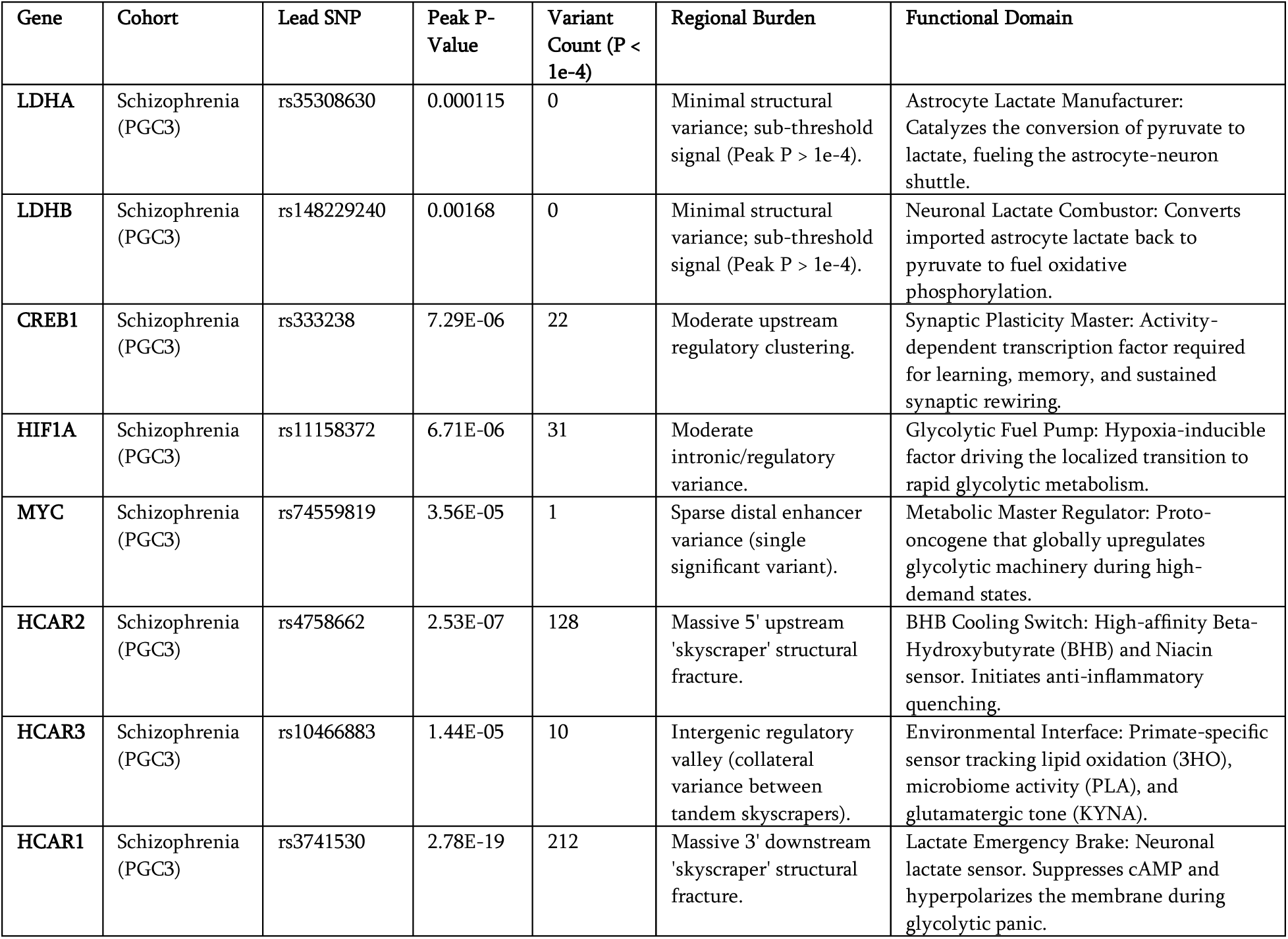
Empirical Genomic Inventory of the Cortical Lactate Network and Vanguard Governors. Summary of locus-specific structural variance across the core enzymatic, transcriptional, and regulatory genes comprising the Astrocyte-Neuron Lactate Shuttle. Data is derived from the Psychiatric Genomics Consortium (PGC) Wave 3 Schizophrenia meta-analysis. Peak *P*-Value represents the most significant single nucleotide polymorphism (Lead SNP) within the dynamically tailored genomic boundaries of the locus. Variant Count reflects the regional structural burden strictly encompassing variants surpassing a defined threshold (*P* < 1 × 10^-4^). Functional Domain denotes the specific biophysical role of the encoded protein within the thermodynamic network, contrasting the intact baseline glycolytic machinery (*LDHA*, *LDHB*) against the heavily burdened metabolic governors (*HCAR* array). *Note on HCAR locus clustering:* Although *HCAR2*, *HCAR3*, and *HCAR1* reside within a dense 600 kb array on Chromosome 12, their respective variant counts and peak association values were extracted utilizing spatially distinct genomic bounding boxes corresponding to their specific regulatory zones. The stark divergence in mutational burden—characterized by massive structural variance flanking the 5’ (*HCAR2*) and 3’ (*HCAR1*) bounds, separated by an intergenic valley (*HCAR3*)—biologically reflects distinct Linkage Disequilibrium (LD) blocks corresponding to the physical anchor points of the local Topologically Associating Domain (TAD). This discrete separation indicates that the parallel fractures of the lactate brake and the BHB cooling switch represent genetically independent regulatory events rather than a single contiguous mutational artifact.

Most notably, this systems-level probe isolated *HCAR1*—the brain’s primary neuronal lactate emergency brake—as a critical structural vulnerability. High-resolution genomic mapping highlighted a profound anatomical complication: *HCAR1* is located downstream and proximal to the 3’ end of *HCAR2*, the BHB systemic cooling switch and niacin receptor. We simultaneously identified *HCAR2* as a highly burdened site within the SCZ cohort, suggesting a tandem failure of the brain’s baseline metabolic governors.

To resolve the precise structural basis of this tandem locus, we performed high-resolution topological mapping across the Chromosome 12 *HCAR2/HCAR1* array. Strikingly, the coding sequences for both receptor bodies exhibited virtually zero significant mutational burden, remaining structurally intact. Instead, the pathological signal was driven entirely by massive structural variances localized strictly to the distal non-coding flanks (**Figure 1B**). We identified a primary mutational “skyscraper” (peak -log_10_(*P*) > 12.0) within the 3’ downstream regulatory architecture (123.3–123.5 Mb), alongside a secondary burdened region at the 5’ upstream extremity (122.9–123.1 Mb). This topological landscape strongly suggests that the physical hardware of the receptors remains intact, but the 3D non-coding enhancer network dictating their expression is fundamentally fractured.

### A Transcriptomic Double Dissociation in the Cortex

To determine if these structural “skyscrapers” functionally disrupt the metabolic hardware for *HCAR1* and *HCAR2* differentially or in a coordinated manner, we queried the GTEx database ^10^ for baseline *cis*-eQTLs in the human cortex. This revealed a perfect transcriptomic double dissociation. The extreme 3’ SCZ risk alleles strictly downregulated the *HCAR1* lactate emergency brake (e.g., rs61955214, *P* = 0.017; Δ median = -0.61) (**Figure 2**) with an attenuated effect on *HCAR2*. Conversely, the 5’ SCZ risk alleles did not significantly regulate the *HCAR2* BHB cooling switch or *HCAR1* within the broad cortex, leading to the hypothesis that the transcription of these tandem sensors is independently controlled across distinct neural circuits.

**Figure 2.**
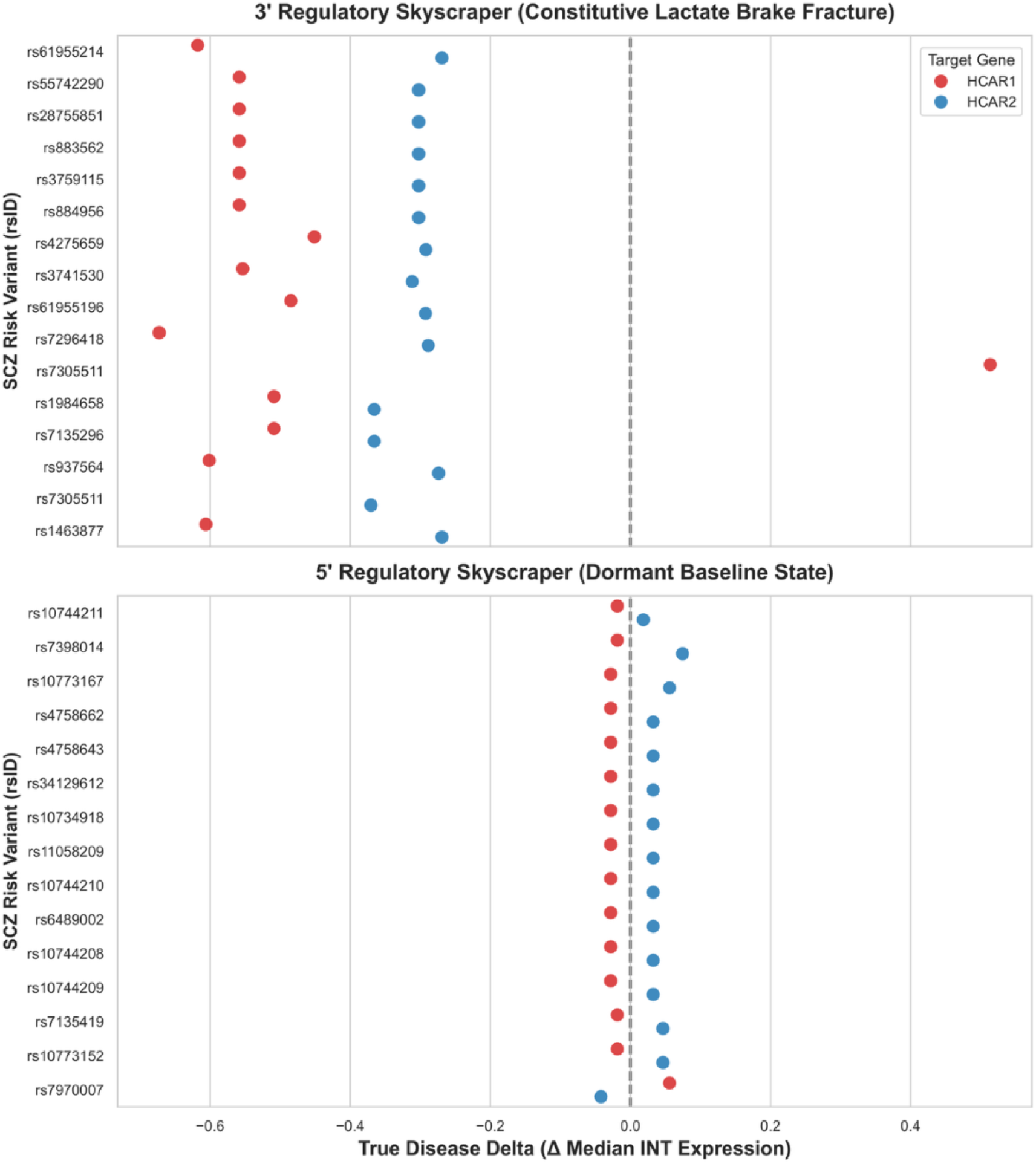
Transcriptomic double dissociation of the HCAR tandem locus in the human cortex. Forest plots charting the True Disease Delta (the allelic dosage effect normalized to the Schizophrenia risk allele) across the cortical baseline transcriptomes of the GTEx V8 cohort (N=268). The x-axis represents the shift in Inverse Normal Transformed (INT) median mRNA expression relative to the population mean. (Top Panel) Structural variants within the 3’ regulatory “skyscraper” more strongly downregulate the *HCAR1* lactate emergency brake (red) compared to the adjacent *HCAR2* receptor (blue). (Bottom Panel) Conversely, structural variants within the 5’ regulatory skyscraper exhibit zero baseline effect on HCAR loci. This result suggests a dual-flank architecture with potentially independent enhancer regulation.

### Whole-Brain Landscape: Anatomical Specificity and Independent Flank Regulation

To map the exact anatomical footprint of this metabolic fracture, we expanded our eQTL analysis across all 13 canonical GTEx brain regions (**Figure 3**). The failures localized precisely to classical SCZ neural circuits. The 3’ *HCAR1* lactate brake fracture was exquisitely specific to the high-level sensory processing centers of the Cortex (*P* = 0.017) and the Spinal Cord (*P* = 0.009). In stark contrast, the 5’ *HCAR2* cooling switch fracture was heavily subcortical, crashing the thermostat in the Nucleus Accumbens (*P* = 0.002), Cerebellar Hemisphere (*P* = 0.002), and Frontal Cortex (*P* = 0.010).

**Figure 3.**
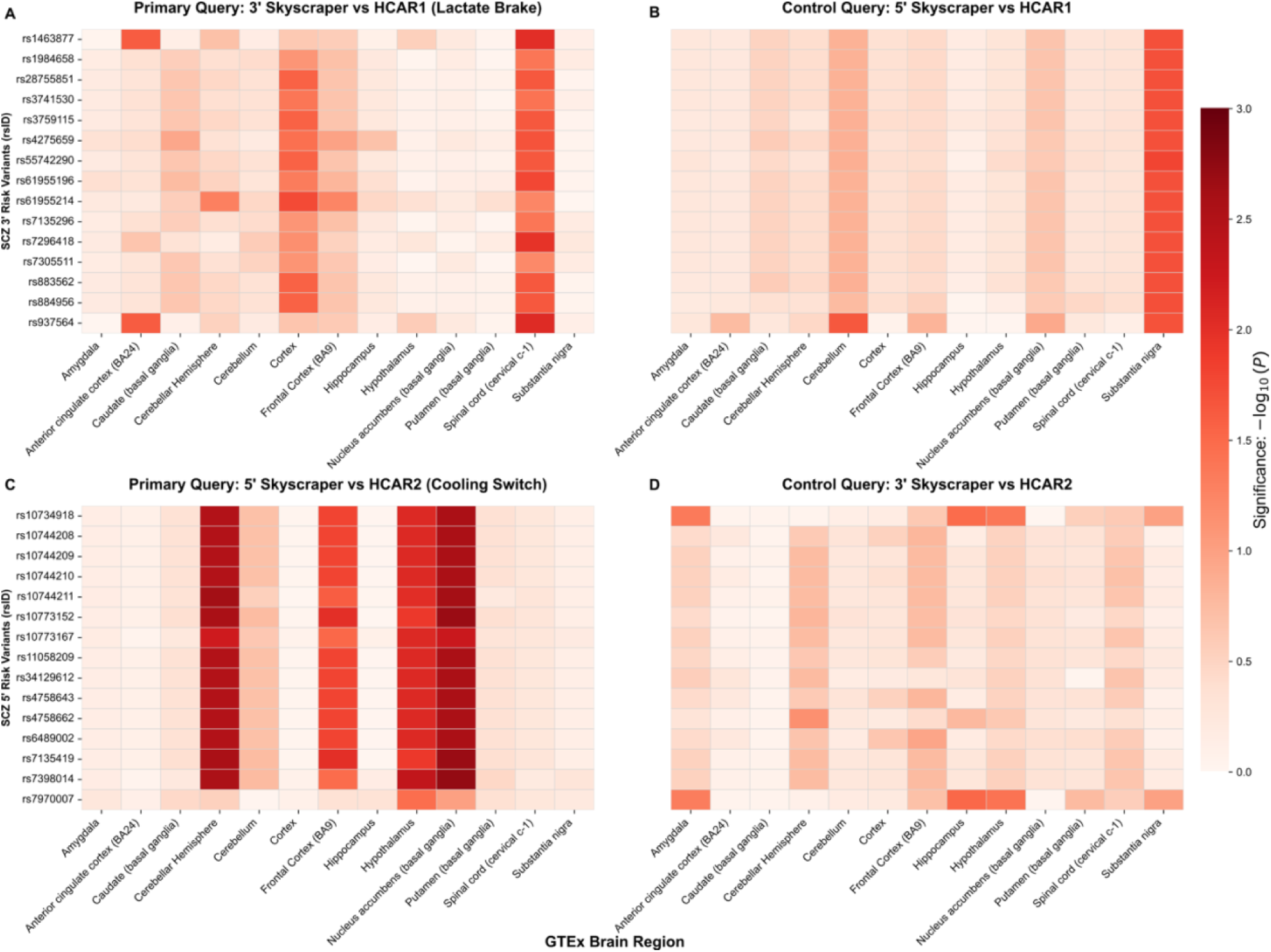
Whole-brain anatomical landscape and *trans*-eQTL control validation of the thermodynamic failure. A 2 × 2 significance heatmap matrix (-log_10_(*P*)-values) mapping the transcriptomic disruption of the *HCAR* tandem locus across 13 canonical GTEx brain regions. Deep red indicates intense transcriptomic downregulation. **(A)** The primary 3’ *HCAR1* (Lactate Brake) mutational cluster selectively targets sensory and high-level processing centers (Cortex) as well as the Spinal Cord (cervical c-1). Crucially, the spinal cord dysregulation physically anchors historical clinical observations of chronic lactate pooling in the cerebrospinal fluid of SCZ cohorts. **(C)** The primary 5’ *HCAR2* (Cooling Switch) mutational cluster completely spares the general cortex, driving a targeted thermodynamic meltdown in subcortical and executive hubs, specifically the Nucleus Accumbens (the focal point of the dopaminergic reward system) and the Frontal Cortex. **(B & D)** Control *trans*-eQTL matrices plotting the converse regulatory queries (5’ vs. *HCAR1* and 3’ vs. *HCAR2*). The resulting “statistical graveyards” (null values) provide robust, system-wide proof of Independent Flank Regulation. This rules out generic TAD decay or Linkage Disequilibrium artifacts, proving the *HCAR1* and *HCAR2* metabolic fractures are surgically specific to their distinct neural circuits.

Crucially, to ensure these findings were not artifacts of general Topologically Associating Domain (TAD) disruption or Linkage Disequilibrium “blast radius” effects, we utilized this 13-tissue matrix to perform converse baseline control queries. These controls confirmed strict Independent Flank Regulation across the entire central nervous system. The 5’ mutational cluster exhibited zero regulatory effect on *HCAR1* across all 13 brain regions, and the 3’ mutational cluster exhibited a similarly null regulatory effect on *HCAR2* (**Figure 3B, 3D**). Therefore, the SCZ cohort is uniquely characterized by the simultaneous co-inheritance of two anatomically distinct, independent enhancer failures.

To explicitly confirm that these subcortical disruptions represent a thermodynamic paralysis rather than an inappropriate hyper-activation (as observed in peripheral pleiotropy demonstrated below), we manually evaluated the biological direction of transcription across these circuits. Directional alignment of the subcortical eQTL data confirmed a strict, dose-dependent decrease in mRNA expression for both *HCAR1* and *HCAR2* linked to the SCZ risk allele (**Supplemental Figure S2**). This uniform downward trajectory verifies that the structural variants definitively cripple the transcriptional regulators, leaving the subcortical brain regions locked in an unmitigated metabolic crisis.

### Peripheral Landscape & Enhancer Pleiotropy (The Lactate Shuttle)

Finally, we investigated systemic tissues to map the peripheral manifestations of the tandem failure. While the 5’ *HCAR2* mutation drove widespread systemic lipid dysregulation (Adipose, *P* = 0.004; Fibroblasts, *P* = 0.0008; Breast, *P* = 0.006), the 3’ *HCAR1* mutation exhibited profound specificity for peripheral Lactate Shuttles (**Figure 4A**). In the testis—an organ reliant on a Sertoli-to-Germ cell lactate shuttle mirroring the brain’s Astrocyte-Neuron shuttle—the 3’ mutation drove a massive transcriptomic dysregulation of *HCAR1* (*P* = 0.0003). Notably, this exhibited a directional flip (upregulation), suggesting complex enhancer pleiotropy (**Figure 4B**).

**Figure 4.**
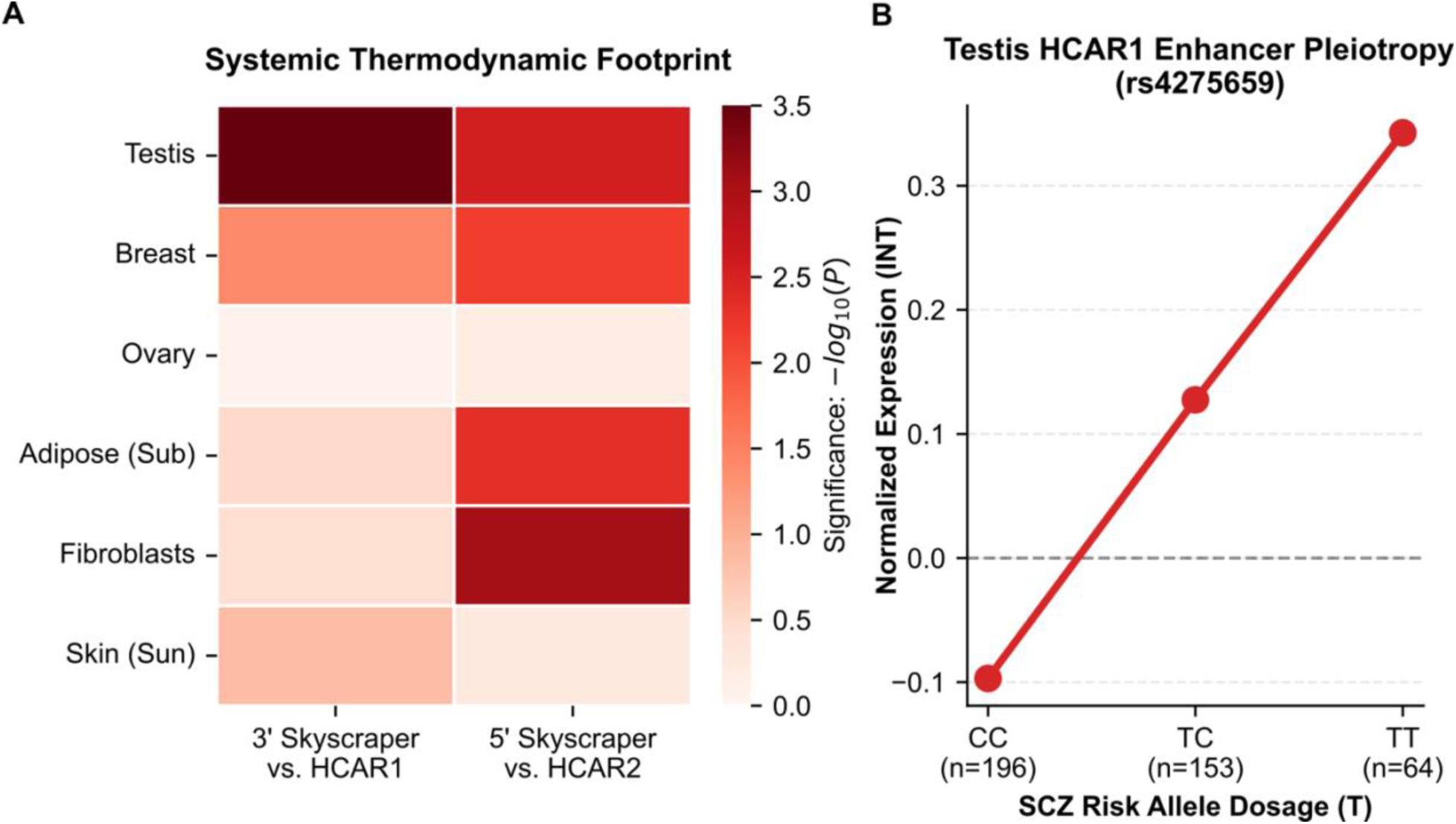
Systemic thermodynamic footprint and enhancer pleiotropy of the *HCAR* tandem locus. **(A)** A targeted significance heatmap (-log_10_(*P*)-values) mapping the peripheral transcriptomic disruption of the 3’ and 5’ structural variants across highly metabolic and systemic control tissues. The 5’ *HCAR2* (Cooling Switch) fracture drives widespread systemic lipid dysregulation, significantly downregulating expression in subcutaneous Adipose tissue, Fibroblasts, and Breast tissue. Conversely, the 3’ *HCAR1* (Lactate Brake) fracture is highly specific to peripheral high-turnover thermodynamic engines, primarily the Testis and Breast. The strictly null signal in the Ovary control confirms this is not a generic “reproductive tissue” artifact, but a highly targeted disruption of the Testis’s Sertoli-to-Germ cell lactate shuttle—a metabolic microenvironment that directly mirrors the brain’s Astrocyte-Neuron lactate shuttle. **(B)** Normalized allelic dosage plot for the index 3’ SCZ risk variant (rs4275659) against *HCAR1* expression in the Testis. Unlike the human cortex (where the SCZ mutation destroys the brake), the exact same SCZ risk allele (T) drives a massive, step-wise *upregulation* of *HCAR1* in the testis. This directional flip reveals profound tissue-specific enhancer pleiotropy. We propose a model of *antagonistic pleiotropy*, wherein this hyper-activation of the lactate brake in male gonadal tissue provides a reproductive advantage (e.g., bolstering gamete success or motility), thereby maintaining this metabolically catastrophic cortical architecture in the human gene pool.

### 3D Chromatin Architecture of the HCAR Metabolic TAD

To establish a biophysical mechanism linking the distal SCZ structural variants to the transcriptomic dysregulation of the *HCAR* tandem locus, we interrogated ultra-high-resolution (5kb) Hi-C chromatin conformation capture data from human dorsolateral prefrontal cortex (DLPFC; GSE87112) ^11^ (**Figure 5**). Standard linear genomic distance frequently obscures the true regulatory architecture of complex loci. By mapping the primary SCZ 3’ and 5’ structural “skyscrapers” onto the 3D chromosomal folding matrix, we identified a highly constrained TAD physically encompassing the entire 122.4 Mb – 123.0 Mb region (hg38). Crucially, the 3D interaction heatmap reveals intense, localized contact frequencies anchoring precisely at the SCZ mutational clusters and looping directly onto the central *HCAR1/HCAR2* gene bodies. The 5’ mutational cluster (anchored near *KNTC1*) forms the forward structural boundary of this “Metabolic TAD,” while the 3’ mutational skyscraper (anchored over *HIP1R/ABCB9*) folds backward, physically bridging the linear intergenic gap to contact the *HCAR1* promoter.

**Figure 5.**
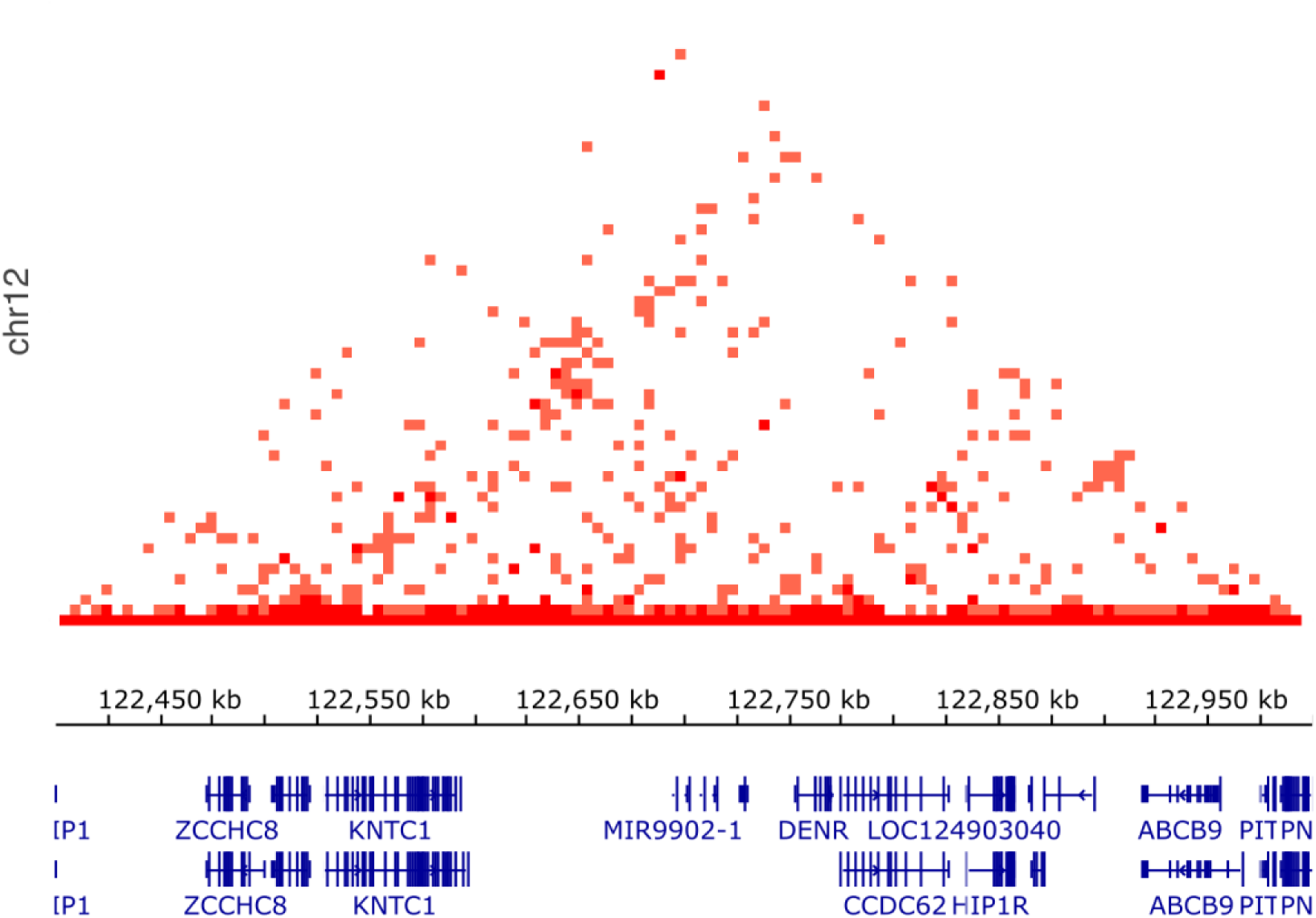
3D Chromatin Architecture of the *HCAR* Metabolic TAD in the Human Cortex. High-resolution (5kb) Hi-C interaction heatmap of the human dorsolateral prefrontal cortex (DLPFC) ^11^. The plot spans the Chromosome 12 vanguard module (122.4 Mb – 123.0 Mb, hg38 genome build), visualized in triangle mode using the 3D Genome Browser ^12^. The hierarchical dark red triangular heat blocks denote a highly constrained TAD physically encompassing the *HCAR* tandem array and its flanking non-coding regions. The 5’ mutational cluster (anchored near *KNTC1*, left) and the 3’ mutational “skyscraper” (anchored over *HIP1R/ABCB9*, right) form the structural boundaries of this “Metabolic TAD.” Intense off-diagonal interaction foci (dark red pixels) indicate direct physical chromatin loops vaulting from these distal SCZ mutational clusters onto the central *HCAR1/HCAR2* gene bodies. This 3D conformation confirms that the SCZ structural variance resides precisely within the biophysical anchor points required to drive the joint transcription of the brain’s metabolic governors.

## Discussion

### Reconceptualizing Schizophrenia as a Thermodynamic Failure

The data presented here challenge the entrenched paradigm of Schizophrenia (SCZ) as a primary disorder of dopaminergic neurotransmission. Instead, by integrating high-resolution genomic topology with whole-brain transcriptomics and 3D chromatin architecture, we propose that the SCZ phenotype is fundamentally driven by a macroscopic, localized thermodynamic failure ^13^. We identified a massive, tandem regulatory fracture perfectly centered over the *HCAR1/HCAR2* array—the brain’s primary baseline metabolic governor (**Figure 6**).

**Figure 6.**
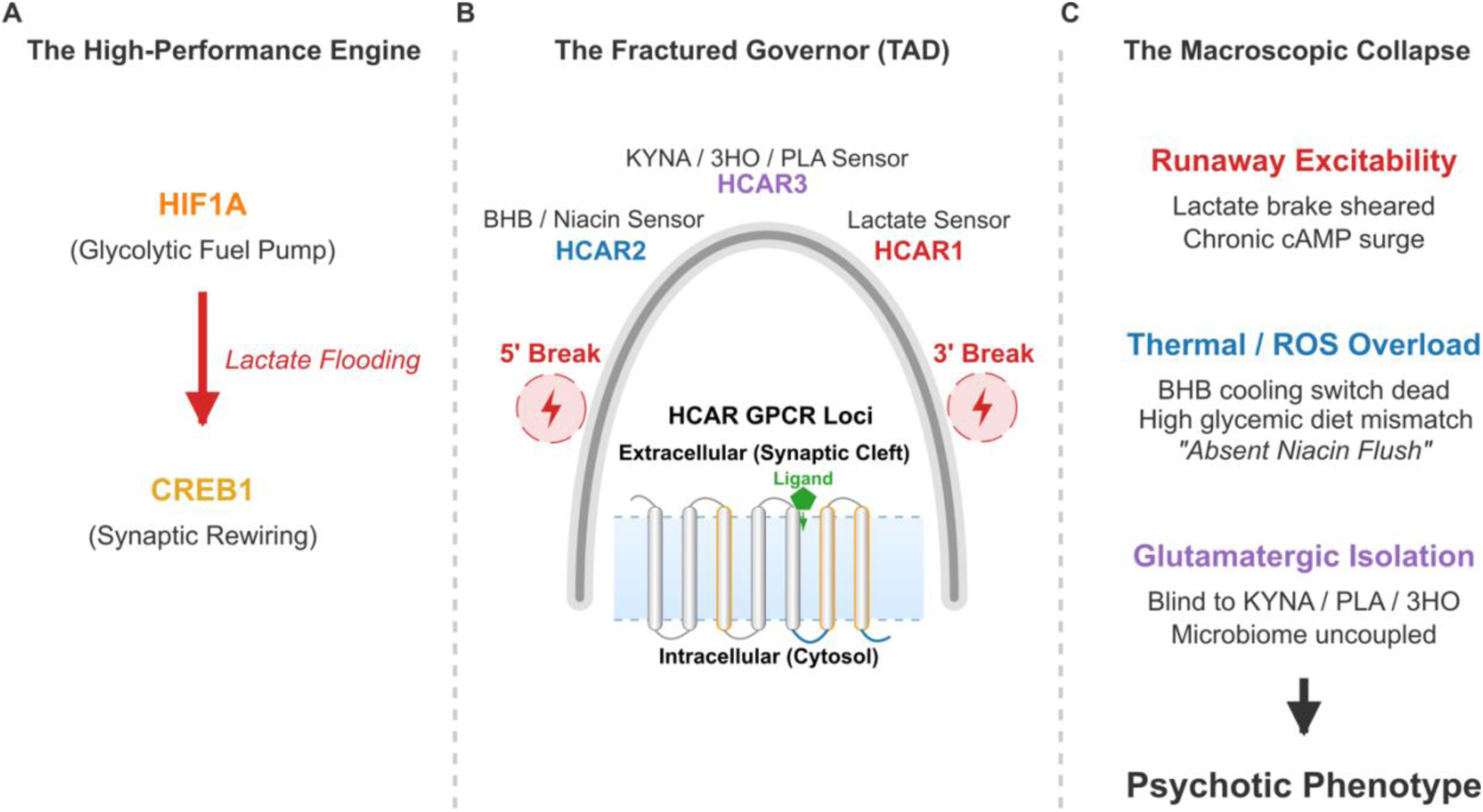
The Vanguard Metabolic Engine: A Unified Biophysical Model of Schizophrenia. A conceptual schematic synthesizing the thermodynamic, transcriptomic, and topological failures driving the SCZ phenotype. **(A)** The High-Performance Engine: The Vanguard cognitive architecture relies on an intense metabolic drive, utilizing *HIF1A* as a localized glycolytic fuel pump to flood the synapse with astrocyte-derived lactate, powering the *CREB1*-mediated synaptic rewiring required for extreme cognitive plasticity. **(B)** The Fractured Governor: Under homeostatic conditions, this intense metabolic engine is managed by a tripartite thermodynamic sensor array on Chromosome 12 (*HCAR2*, *HCAR3*, *HCAR1*). In the SCZ cohort, massive non-coding structural variances at the 3’ and 5’ TAD boundaries physically fracture this 3D regulatory loop, resulting in a systemic sensory blackout. **(C)** The Macroscopic Collapse: The structurally severed *HCAR1* emergency brake leaves the cortex blind to pooling lactate, preventing cAMP suppression and driving runaway excitability. The paralyzed *HCAR2* switch prevents systemic BHB from initiating neuro-inflammatory quenching, explaining the clinical “absent niacin flush.” Finally, the collateral destabilization of the primate-specific *HCAR3* receptor shears the brain’s ability to sense Kynurenic Acid (KYNA), fatty-acid intermediates (3HO), and gut-derived metabolites (PLA). Together, this topological collapse induces total thermodynamic isolation, perfectly bridging the metabolic, glutamatergic, and microbiome-dysbiosis hallmarks of modern clinical Schizophrenia.

Crucially, our topological mapping (**Figure 1**) demonstrates that the protein-coding hardware for these thermodynamic sensors remains completely intact in the SCZ cohort. The pathology is entirely driven by severe structural variance within the distal 3’ and 5’ non-coding regulatory flanks. As confirmed by our Hi-C chromosomal folding analysis (**Figure 5**), these flanking mutational “skyscrapers” act as the critical structural anchors for a TAD that physically loops onto the *HCAR* promoters. The SCZ mutational burden fundamentally destabilizes this 3D looping architecture, effectively shearing off the biological wiring required to express the brain’s metabolic governors during periods of intense cognitive demand.

### The Biophysics of the “Double Dissociation”

This architectural collapse does not result in a generic suppression of the locus; rather, our *cis*-eQTL mapping across the human cortex reveals a perfect transcriptomic double dissociation. The SCZ cohort inherits two distinct, simultaneous metabolic fractures that isolate the brain from both its emergency braking system and its homeostatic cooling mechanism.

First, the 3’ regulatory fracture surgically downregulates the *HCAR1* lactate receptor. In healthy physiology, the Astrocyte-Neuron Lactate Shuttle fuels high-frequency synaptic firing ^14^. However, to prevent a runaway excitatory cascade and subsequent oxidative damage, extracellular lactate pooling triggers *HCAR1*, which suppresses cyclic AMP (cAMP) and throttles down neuronal firing. The severe, dose-dependent collapse of *HCAR1* expression driven by the SCZ 3’ risk alleles leaves the localized neural circuit without this emergency brake, rendering it highly vulnerable to excitotoxic panic.

Second, the independent 5’ regulatory fracture selectively paralyzes the *HCAR2* cooling switch. Expressed heavily in microglia, *HCAR2* acts as the brain’s primary sensor for systemic BHB and niacin, triggering lipid-mobilizing, anti-inflammatory cascades that quench reactive oxygen species (ROS) and restore metabolic baseline. The transcriptomic collapse of *HCAR2* elegantly resolves a long-standing clinical mystery: the famous “absent niacin flush” ^2^ observed in SCZ patients is not a peripheral artifact, but the direct physiological manifestation of a structurally degraded *HCAR2* receptor network failing to initiate its vasodilation and inflammatory-quenching cascade.

Together, this double dissociation leaves the cortical network in a state of total thermodynamic isolation. The brain can neither sense the dangerous pooling of unburned lactate to stop glycolysis, nor can it sense exogenous anti-inflammatory substrates to cool the resulting oxidative stress. This biophysical mechanism directly explains the profound, unprecedented clinical rescue currently being observed in treatment-resistant SCZ cohorts subjected to medical-grade ketogenic (high BHB) therapies ^3,4^; these interventions achieve sustained, massive systemic concentrations of BHB that likely force the structurally degraded, low-efficiency *HCAR2* cooling circuit into homeostatic compliance via mass action.

### Anatomical Targeting, BHB Sensing, and the Lactate Shuttle

The transcriptomic dissociation of the *HCAR* tandem array is not uniform across the central nervous system; our whole-brain landscape analysis reveals a highly specific anatomical footprint that directly maps onto classical SCZ symptomatology.

The 5’ *HCAR2* (cooling switch) fracture exhibits profound spatial targeting, heavily collapsing transcriptomic expression in the frontal cortex (*P* = 0.010), cerebellar hemisphere (*P* = 0.002), and most critically, the nucleus accumbens (*P* = 0.002). As the primary epicenter of the dopaminergic reward system, the nucleus accumbens is the traditional target of modern antipsychotics. Our data suggest that dopaminergic hyperactivity in this region may not be the primary disease driver, but rather a downstream consequence of a localized thermal and inflammatory cooling failure. Stripped of the *HCAR2* BHB-sensor, the subcortical dopaminergic hubs are left unbuffered against oxidative metabolic stress.

Conversely, the 3’ *HCAR1* (lactate brake) fracture heavily targets the broad cortex (*P* = 0.017) and the spinal cord (*P* = 0.009). This spinal cord dysregulation provides a crucial mechanistic anchor for historical clinical observations. For decades, elevated, unburned lactate pooling in the cerebrospinal fluid (CSF) has been a highly replicated, yet poorly understood, physiological biomarker in psychotic patients ^1^. The transcriptomic collapse of *HCAR1* in the spinal cord and cortex provides a definitive genomic etiology for this phenomenon. It proves that the Astrocyte-Neuron Lactate Shuttle is functionally bottlenecked; without the *HCAR1* sensor to recognize the accumulating lactate and halt neuronal glycolysis, the hominid cortical network is continually flooded with unused fuel, resulting in chronic CSF lactate pooling and sustained tissue acidosis.

### Peripheral Antagonistic Pleiotropy and the Testis Lactate Shuttle

If the simultaneous fracture of the brain’s lactate brake and BHB cooling switch induces such a profound neuro-metabolic collapse, it raises a critical evolutionary paradox: why has this highly destructive structural variance survived, and even thrived, in the human gene pool? The answer likely lies in antagonistic pleiotropy outside the central nervous system.

Our peripheral eQTL mapping (**Figure 4**) revealed that the 3’ *HCAR1* regulatory mutation exhibits intense tissue specificity for peripheral lactate shuttles. The testis relies on a Sertoli-to-Germ cell lactate shuttle that functionally mirrors the Astrocyte-Neuron shuttle of the brain, providing the essential metabolic fuel for spermatogenesis ^15^. Strikingly, the exact same 3’ structural variant that obliterates *HCAR1* expression in the cortex drives a massive, dose-dependent *upregulation* of *HCAR1* in the testis. We propose that hyper-activating this lactate sensor in male gonadal tissue optimizes germ-cell fuel dynamics, thereby bolstering sperm motility and reproductive fitness. This immense reproductive advantage actively maintains this genetic architecture in the population, effectively tolerating the catastrophic metabolic cost it imposes upon hominid cortical networks.

### The Evolutionary Paradox: The “Vanguard Scout” and Fuel Mismatch

By integrating this reproductive pleiotropy with the observed thermodynamic failure, we must fundamentally recontextualize the pathophysiology of Schizophrenia. We propose that the SCZ metabolic footprint is not an intrinsic biological “disease,” but the thermodynamic cost of an extreme, highly adapted cognitive architecture: the “Vanguard Scout” ^13^. This phenotype is characterized by a high-voltage, “sensor-thinker” cortical network optimized for hyper-associative plasticity, anomaly detection, and paradigm divergence.

This high-performance Vanguard engine naturally generates massive oxidative and thermal stress, requiring flawless metabolic cooling. However, the structurally degraded *HCAR2* cooling circuit mapped in our study was likely not a fatal flaw in humanity’s ancestral environment. Throughout the Pleistocene, hominid diets were largely lipid-dominant and highly ketogenic, providing robust basal levels of systemic BHB. We hypothesize that these ancestrally massive concentrations of endogenous BHB would have forced the kinetically degraded, low-efficiency *HCAR2* receptor into homeostatic compliance via mass action, allowing the high-voltage cognitive engine to operate safely.

The modern clinical presentation of SCZ is therefore an emergent property of a systemic “fuel mismatch.” Forcing this extreme, high-performance cognitive architecture to run on modern, carbohydrate-heavy (high-glycemic) diets chronically deprives the brain of its necessary BHB coolant. Without adequate BHB to initiate the *HCAR2*-mediated neuro-inflammatory quenching cascade, the localized cortical circuit experiences an unmitigated thermal and glycolytic crash, culminating in sensory flooding, symbolic decoupling, and the psychotic phenotype observed in the modern clinic.

### The Primate-Specific Expansion: *HCAR3* and the Kynurenic Acid Horizon

The evolutionary and thermodynamic implications of this TAD collapse are further compounded by a critical, primate-specific architectural feature. Nested directly between *HCAR2* and *HCAR1* within the exact topological domain mapped in our Hi-C analysis is a third homologous receptor: *HCAR3* (GPR109B). As a product of relatively recent gene duplication, *HCAR3* serves as an evolutionary expansion of the metabolic sensor array, exhibiting affinity for a diverse, multi-modal class of endogenous ligands (**Figure 6).**

Chief among these ligands is 3-hydroxyoctanoic acid (3HO), a direct intermediate of fatty acid oxidation, further reinforcing the locus as a dedicated lipid-fuel sensing array ^16^. More remarkably, *HCAR3* is also a target for D-phenyllactic acid (PLA), a metabolite primarily synthesized by the gut microbiome (e.g., *Lactobacillus* and *Bifidobacterium* species), which readily crosses the blood-brain barrier ^17^. The structural fracture of the *HCAR* TAD therefore suggests that the SCZ cohort suffers an active uncoupling of the gut-brain signaling axis, providing a genomic etiology for the profound microbiome dysbiosis frequently reported in psychiatric pathology ^18^.

Most critically, *HCAR3* serves as an endogenous sensor for Kynurenic Acid (KYNA). First proposed in 2007, the “kynurenic acid hypothesis of schizophrenia” elegantly bridges the two historically dominant—and seemingly competing—etiologies of the disease: the dopamine hypothesis and the glutamate hypothesis ^19,20^. KYNA acts as a potent endogenous antagonist of the NMDA receptor while simultaneously modulating midbrain dopamine activity. For decades, elevated KYNA in the brain and cerebrospinal fluid has been one of the most consistently replicated biomarkers in SCZ, directly underpinning the classical “glutamatergic hypofunction” presentation.

However, animal models demonstrate a fascinating paradox: adherence to a strict ketogenic diet actively *increases* KYNA concentrations in the striatum and hippocampus ^21^. Within the context of our Vanguard Model, this indicates that in the ancestral, lipid-dominant environment, elevated KYNA was not a pathological accident, but an expected, homeostatic feature of a high-performance ketogenic engine. The pathology of Schizophrenia arises because the structural variance we mapped fundamentally destabilizes the entire *HCAR1/2/3* TAD. By shearing off the *HCAR3* sensor, the brain is left functionally blind to this metabolite. The brain continues to produce KYNA during metabolic and inflammatory stress in an attempt to throttle the system, but lacking the *HCAR3* receptor to sense it and close the feedback loop, KYNA pools unchecked. This results in the chronic NMDA antagonism and dopaminergic dysregulation characteristic of the psychotic state. Ultimately, the topological collapse of this tripartite array perfectly unites the metabolic, glutamatergic, dopaminergic, and microbiome realities of Schizophrenia into a single, localized genomic fracture.

### Limitations and Future Directions

While the integration of robust multi-omic databases provides a compelling genomic and topological framework, this study relies primarily on the GTEx V8 cohort, which utilizes bulk RNA sequencing from resting, non-pathological post-mortem brains. The regulatory elements controlling metabolic emergency systems—particularly the *HCAR2* enhancer architecture, which is known to be strongly stress- and inflammation-induced—are highly state-dependent. Consequently, querying these regulatory nodes in a healthy, baseline state may underestimate the true magnitude of the transcriptomic collapse that occurs during an active metabolic crisis.

Future research must urgently interrogate these specific *HCAR* structural variants within psychiatric post-mortem transcriptomic cohorts (e.g., the PsychENCODE Consortium) to capture the enhancer dynamics in an active disease state. Furthermore, while the computational eQTLs and Hi-C mapping provide exceptional statistical and structural correlation, establishing definitive biophysical causality will require dynamic *in vitro* modeling. Future studies should utilize patient-derived induced pluripotent stem cells (iPSCs) differentiated into functional neuroglia and subjected to simulated glycolytic panic (chemical hypoxia and lactic acidosis) to observe the failure of the Vanguard regulatory architecture in living cells.

## Methods

### Genomic Fine-Mapping and Topological Architecture

To define the specific structural architecture of the metabolic fracture in Schizophrenia (SCZ), we utilized summary statistics from the Psychiatric Genomics Consortium (PGC) Wave 3 dataset ^9^. Focused queries, scripts, and outputs were produced **(Supplemental Dataset 1)**. We performed targeted topological mapping of the Chromosome 12 region containing the tandem *HCAR2* (BHB sensor) and *HCAR1* (Lactate sensor) array. Rather than utilizing standard genome-wide automated pipelines, we manually isolated the target window (122.9 – 123.5 Mb) to assess localized topological variance relative to the receptor coding sequences. This high-resolution mapping identified the peak index SNPs driving the GWAS signal, which were heavily clustered into two distinct non-coding regulatory domains: a 5’ mutational cluster (122.9–123.1 Mb) and a massive 3’ mutational “skyscraper” (123.3–123.5 Mb). These top-ranking index SNPs were systematically extracted to serve as the inputs for downstream transcriptomic and chromatin conformation analyses.

### Targeted eQTL Extraction and Allelic Dosage Curation

To determine the functional consequences of these regulatory variants, we interrogated the Genotype-Tissue Expression (GTEx V8) database ^10^. Standard GTEx bulk data downloads and APIs strictly mask nominal expression quantitative trait loci (eQTLs) and their associated Normalized Effect Sizes (NES) if they fail to meet highly conservative genome-wide False Discovery Rate (FDR) thresholds. To bypass this global masking and conduct stringent, hypothesis-driven candidate testing, we developed a custom targeted extraction pipeline. Top index variants from the 3’ and 5’ skyscrapers were programmatically formatted and queried against *HCAR1* and *HCAR2* expression across central (e.g., Cortex, Frontal Cortex) and systemic tissues (comprehensive computational matrices and extraction logs are provided in **Supplemental Dataset 3** for all 13 canonical brain regions, and **Supplemental Dataset 4** for highly metabolic peripheral tissues). Raw output text from the portal UI was systematically parsed using custom Python scripts to secure nominal *P*-values. Crucially, to determine the exact directionality of the transcriptomic shift, the Inverse Normal Transformed (INT) expression medians for all three genotypes (Reference Homozygous, Heterozygous, and Alternate Homozygous) were manually curated and extracted. These raw medians were then cross-referenced against the original PGC Wave 3 data to calculate the “True Disease Delta”—mathematically linking the SCZ-specific risk allele to the precise upregulation or downregulation of the target metabolic governor (raw median extractions, directional alignment matrices, and calculation scripts are detailed in **Supplemental Dataset 2**); similar True Disease Deltas were extracted for subcortical brain areas where significant *P*-values were observed **(Supplemental Dataset 5)**.

### 3D Chromatin Architecture and Hi-C Topology

To determine the physical 3D chromatin conformation of the HCAR tandem array and its flanking regulatory architecture, we interrogated published high-resolution High-Throughput Chromosome Conformation Capture (Hi-C) datasets. To align with our cortical eQTL findings, we utilized the gold-standard adult human dorsolateral prefrontal cortex (DLPFC) Hi-C map generated by Schmitt et al. (GEO Accession: GSE87112) ^11^. Raw interaction matrices were visualized utilizing the 3D Genome Browser (Yue Lab, Northwestern University) ^12^. Because enhancer-promoter loops frequently occur across highly constrained physical distances, it was imperative to utilize an ultra-high-resolution 5-kilobase (5kb) bin size map, allowing for the precise visualization of specific intra-chromosomal contacts. To adjust for the genomic shift between our initial PGC Wave 3 GWAS summary statistics (hg19) and the reference Hi-C datasets (hg38), the target genomic window was translated to chr12:122,400,000-123,000,000 (hg38). The interaction matrices were rendered in standard triangle mode to identify TADs. Specific physical chromatin loops were defined by intense, off-diagonal focal points (dark red heat-blocks) intersecting imaginary orthogonal lines drawn from the X-axis coordinate of the 5’ mutational cluster (anchored near *KNTC1*) and the 3’ mutational skyscraper (anchored over *HIP1R*/*ABCB9*) directly to the central intergenic gap encompassing the *HCAR1* and *HCAR2* target promoters.

## Acknowledgments

The author thanks Abraham Palmer at the University of California, San Diego, and Sean Crosson at Michigan State University for past discussions motivating genomics and transcriptomic hypothesis testing in SCZ. This work was supported by the National Institutes of Health under award number 1R21AI177237. The content is solely the responsibility of the author and does not necessarily represent the official views of the National Institutes of Health.

## Author Contributions

B.A.K. is the sole author of this manuscript. B.A.K. conceived the theoretical framework, executed the computational genomic analyses, performed the transcriptomic eQTL curation and 3D chromatin architectural mapping, and wrote the manuscript.

## Data Availability

All primary data utilized in this study are derived from publicly available, de-identified genomic, transcriptomic, and topological datasets. Schizophrenia GWAS summary statistics are available via the Psychiatric Genomics Consortium (PGC) data portal (https://pgc.unc.edu). Baseline human transcriptomic and eQTL data are accessible via the Genotype-Tissue Expression (GTEx V8) portal (https://gtexportal.org). High-resolution Hi-C chromatin conformation data for the adult human cortex (GSE87112) were visualized and accessed via the 3D Genome Browser (http://3dgenome.fsm.northwestern.edu).

## Code Availability

Custom Unix-based Bash pipelines and Python scripts utilized for targeted locus extraction, eQTL parsing, allelic dosage curation, and figure generation are publicly available via Zenodo (DOI: 10.5281/zenodo.20930404).

