## Supplemental Figures Document for "State-Dependent Transcriptomic Collapse of the Brain’s Lactate and Ketone Thermodynamic Sensors in Schizophrenia"

Bryan A. Krantz

Department of Microbial Pathogenesis, School of Dentistry, University of Maryland, Baltimore, 650 W. Baltimore Street, Baltimore, MD 21201, U.S.A.

 (BAK)

**Running title:** Thermodynamic Sensors in Schizophrenia

**Keywords:** Metabolic Psychiatry, Schizophrenia, Lactate, Beta-Hydroxybutyrate (BHB), Genomics, Transcriptomics, eQTL, HCAR1 (GPR81), HCAR2 (GPR109A), HCAR3 (GPR109B), Astrocyte-Neuron Lactate Shuttle

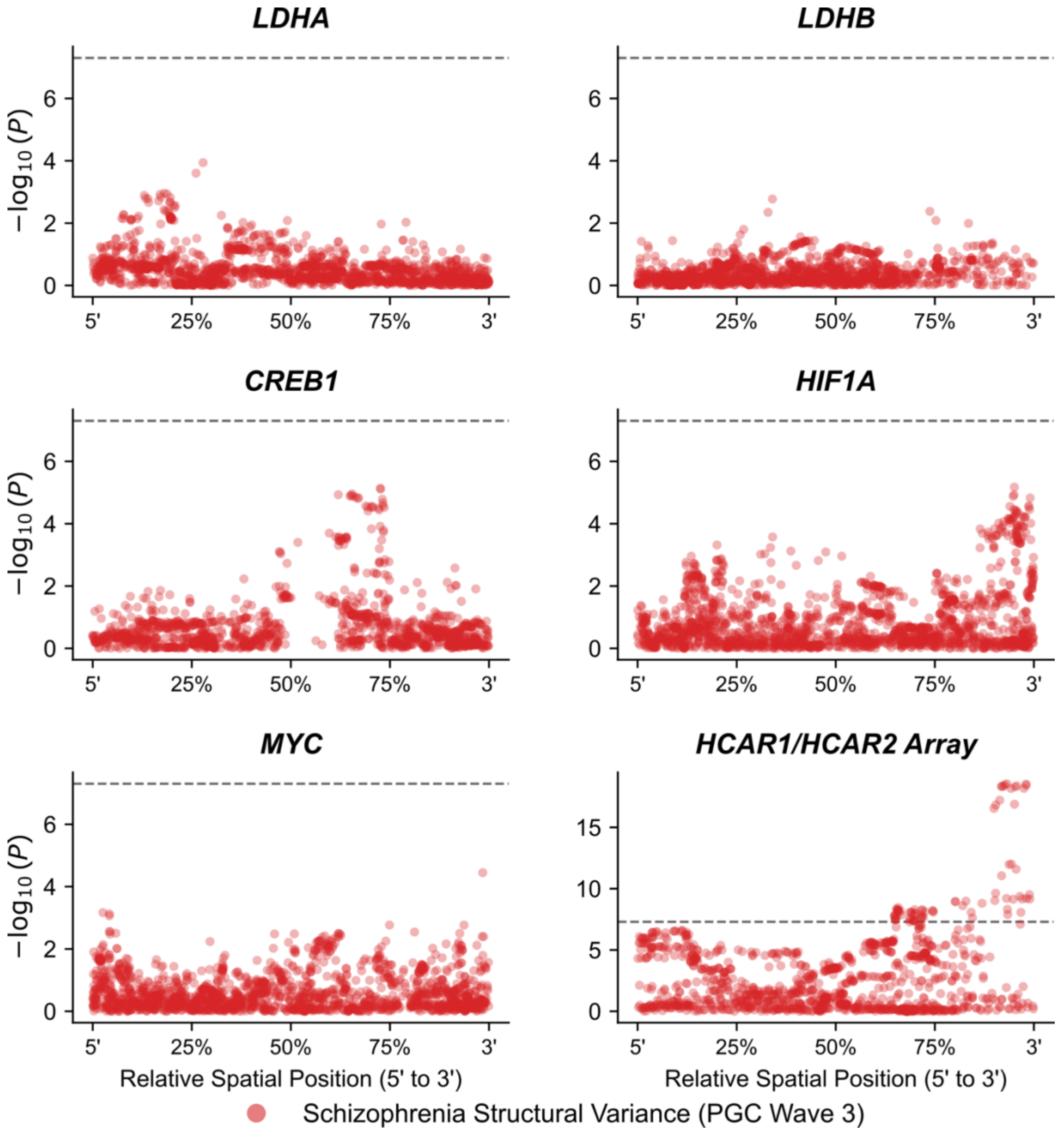


**Supplemental Figure S1. Regional Genomic Burden of the Cortical Lactate Network in Schizophrenia.** A multi-panel locus-zoom grid detailing the dense structural variance across the six core loci comprising the astrocyte-neuron lactate shuttle and the "Vanguard" metabolic engine. Data is derived from the Psychiatric Genomics Consortium (PGC) Wave 3 Schizophrenia meta-analysis. To visualize discrete genomic windows across disparate chromosomes on a single standardized scale, absolute base-pair coordinates were geometrically normalized. The x-axis represents the relative spatial position from the 5' upstream regulatory boundary to the 3' downstream regulatory boundary of each specific extraction window. The y-axis denotes statistical association ($-\log_{10}(P)$), with the dashed black line indicating standard genome-wide significance ($P = 5 \times 10^{-8}$). The network highlights significant regional vulnerability across the distributed metabolic infrastructure, including: the astrocyte lactate manufacturer (*LDHA*), the neuronal lactate combustor (*LDHB*), the master regulator of activity-dependent synaptic plasticity (*CREB1*), the hypoxic/glycolytic fuel pump (*HIF1A*), the glycolytic master regulator (*MYC*), and the tandem thermodynamic governor array (*HCAR1/HCAR2*). This network-wide footprint demonstrates that the polygenic architecture of Schizophrenia heavily burdens the coordinated enzymatic and transcriptional nodes required to maintain high-voltage cortical metabolism.

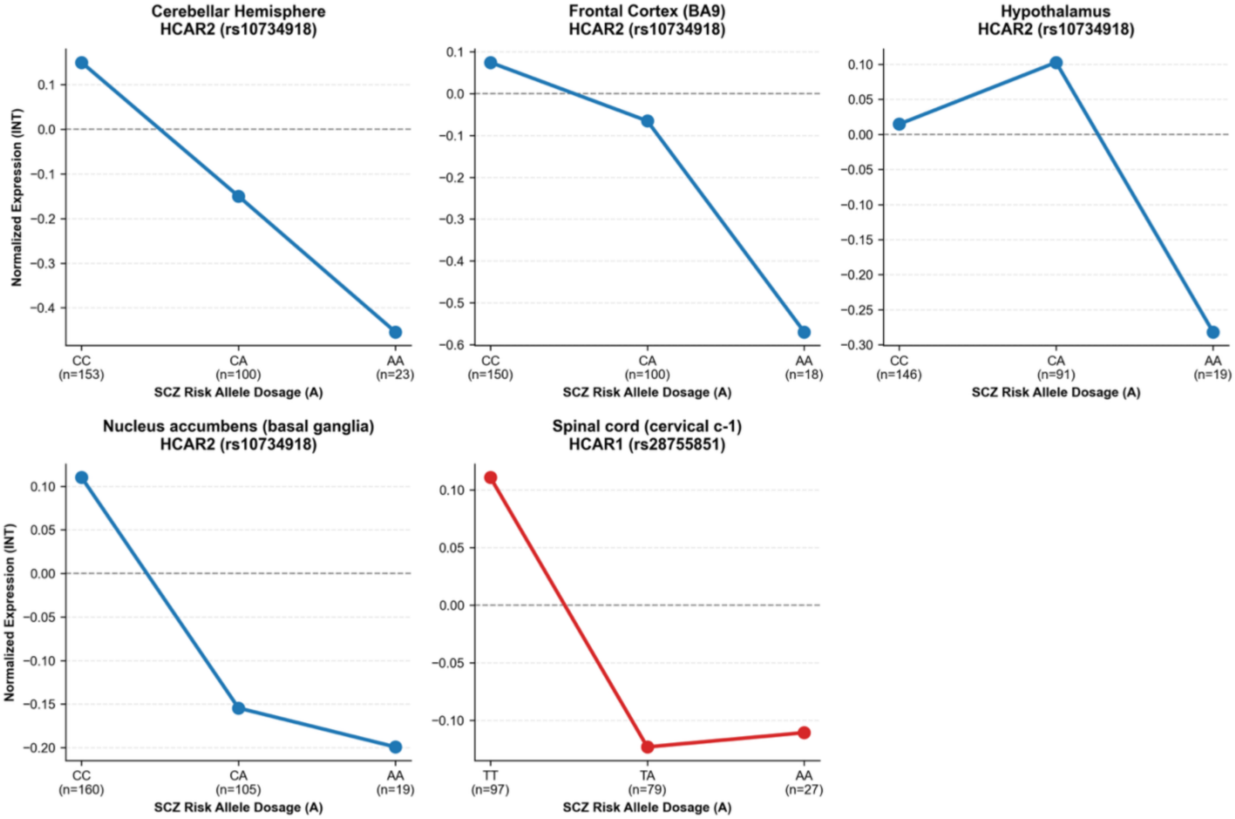


**Supplemental Figure S2. Directional Transcriptomic Collapse of the *HCAR* Governors Across Subcortical Neural Circuits.** A multi-panel dosage grid plotting the Inverse Normal Transformed (INT) median mRNA expression against the Schizophrenia (SCZ) risk allele dosage (0, 1, or 2 copies). Manual extraction and directional alignment of GTEx V8 subcortical eQTL data confirm a strict, dose-dependent downregulation of both the *HCAR1* lactate brake (e.g., Spinal Cord) and the *HCAR2* cooling switch (e.g., Nucleus Accumbens, Cerebellar Hemisphere, Hypothalamus). The uniform downward trajectory across these tissues confirms that the SCZ mutational burden induces a systemic paralysis of the brain's metabolic governors, explicitly ruling out intra-brain enhancer pleiotropy.
