## Supplemental Dataset 1 for "State-Dependent Transcriptomic Collapse of the Brain’s Lactate and Ketone Thermodynamic Sensors in Schizophrenia": bash-lactate-series-SCZ-PGC.sh

```
# 1. Extract the Astrocyte Manufacturer (LDHA - Chromosome 11)
zgrep -v "^##" PGC3_SCZ_wave3.primary.autosome.public.v3.vcf.tsv.gz | awk 'NR==1 {print} $1=="11" && $3>=18090000 && $3<=18710000 {print}' > Lactate_LDHA_wide_regulatory.tsv

# 2. Extract the Neuronal Combustor (LDHB - Chromosome 12)
zgrep -v "^##" PGC3_SCZ_wave3.primary.autosome.public.v3.vcf.tsv.gz | awk 'NR==1 {print} $1=="12" && $3>=21340000 && $3<=21970000 {print}' > Lactate_LDHB_wide_regulatory.tsv

# 3. Major Lactate Sensor GPCR
zgrep -v "^##" PGC3_SCZ_wave3.primary.autosome.public.v3.vcf.tsv.gz | awk 'NR==1 {print} $1=="12" && $3>=122910000 && $3<=123515000 {print}' > Lactate_HCAR1_wide_regulatory.tsv

# 4. Wide Regulatory Sweep of HCAR1 and HCAR2 Loci
zgrep -v "^##" PGC3_SCZ_wave3.primary.autosome.public.v3.vcf.tsv.gz | awk 'NR==1 {print} $1=="12" && $3>=122850000 && $3<=123515000 {print}' > Tandem_HCAR2_HCAR1_wide_regulatory.tsv
```
