## Supplemental Dataset 1 for "State-Dependent Transcriptomic Collapse of the Brain’s Lactate and Ketone Thermodynamic Sensors in Schizophrenia": bash-lactate-transcriptional-regulatory-series-SCZ-PGC.sh

```
# 1. The Synaptic Plasticity Master (CREB1 - Chromosome 2) (hg19 window: ~108.11 Mb to 108.79 Mb)
zgrep -v "^##" PGC3_SCZ_wave3.primary.autosome.public.v3.vcf.tsv.gz | awk 'NR==1 {print} $1=="2" && $3>=108110000 && $3<=108790000 {print}' > Transcript_CREB1_wide_regulatory.tsv

# 2. The Hypoxic/Metabolic Stress Responder (HIF1A - Chromosome 14) (hg19 window: ~61.86 Mb to 62.51 Mb)
zgrep -v "^##" PGC3_SCZ_wave3.primary.autosome.public.v3.vcf.tsv.gz | awk 'NR==1 {print} $1=="14" && $3>=61860000 && $3<=62510000 {print}' > Transcript_HIF1A_wide_regulatory.tsv

# 3. The Glycolytic Master Regulator (MYC - Chromosome 8) (hg19 window: ~128.44 Mb to 129.05 Mb)
zgrep -v "^##" PGC3_SCZ_wave3.primary.autosome.public.v3.vcf.tsv.gz | awk 'NR==1 {print} $1=="8" && $3>=128440000 && $3<=129050000 {print}' > Transcript_MYC_wide_regulatory.tsv
```
