## Supplemental Dataset 1 for "State-Dependent Transcriptomic Collapse of the Brain’s Lactate and Ketone Thermodynamic Sensors in Schizophrenia": README.docx

**Supplemental Dataset 1: Genomic Extraction of the Cortical Lactate Network**

**Study:** State-Dependent Transcriptomic Collapse of the Brain's Lactate and Ketone Thermodynamic Sensors in Schizophrenia

**Author:** Bryan A. Krantz

**Overview**

This repository contains the raw computational extraction scripts and the resulting targeted genomic summary statistics utilized to map the regional mutational burden of the "Vanguard" metabolic engine. The data specifically isolates structural variance across the core enzymatic, transcriptional, and regulatory genes comprising the Astrocyte-Neuron Lactate Shuttle and the *HCAR* thermodynamic governor array.

These .tsv files serve as the direct empirical inputs for generating **Figure 1**, **Supplemental Figure S1**, and **Table 1** in the manuscript.

**Primary Data Source**

All targeted extractions were performed against the **Psychiatric Genomics Consortium (PGC) Wave 3 Schizophrenia Meta-Analysis** summary statistics (build hg19):

- PGC3_SCZ_wave3.primary.autosome.public.v3.vcf.tsv.gz
- Available via: https://pgc.unc.edu

**File Manifest**

**1. Extraction Scripts (Bash)**

These custom Unix-based pipelines utilize zgrep and awk to precisely slice the massive PGC3 GWAS dataset into discrete functional genomic windows.

- bash-lactate-series-SCZ-PGC.sh: Contains the coordinate boundaries and extraction logic for the core lactate machinery (*LDHA*, *LDHB*) and the *HCAR* receptor arrays.
- bash-lactate-transcriptional-regulatory-series-SCZ-PGC.sh: Contains the coordinate boundaries and extraction logic for the upstream metabolic and plasticity transcription factors (*CREB1*, *HIF1A*, *MYC*).

**2. Genomic Output Files (.tsv)**

The resulting targeted summary statistic files. Each file contains all evaluated SNPs within the explicitly defined regulatory boundaries of the target gene, retaining standard GWAS columns (CHROM, POS, ID, PVAL, etc.).

**The Thermodynamic Governor Array:**

- Tandem_HCAR2_HCAR1_wide_regulatory.tsv: The primary locus of the study. A massive wide-regulatory sweep (~600 kb) capturing the 5' and 3' structural "skyscrapers" flanking the *HCAR2* (BHB/Niacin sensor) and *HCAR1* (Lactate sensor) tandem array on Chromosome 12.
- Lactate_HCAR1_wide_regulatory.tsv: A targeted sub-extraction specifically centering on the *HCAR1* (lactate emergency brake) downstream regulatory domain.

**The Baseline Lactate Machinery:**

- Lactate_LDHA_wide_regulatory.tsv: Captures the *LDHA* locus (Chromosome 11) representing the astrocyte lactate manufacturer.
- Lactate_LDHB_wide_regulatory.tsv: Captures the *LDHB* locus (Chromosome 12) representing the neuronal lactate combustor.

**The High-Performance Plasticity & Glycolytic Regulators:**

- Transcript_CREB1_wide_regulatory.tsv: Captures the *CREB1* locus (Chromosome 2), the master regulator of activity-dependent synaptic plasticity and rewiring.
- Transcript_HIF1A_wide_regulatory.tsv: Captures the *HIF1A* locus (Chromosome 14), the hypoxia-inducible glycolytic fuel pump.
- Transcript_MYC_wide_regulatory.tsv: Captures the *MYC* locus (Chromosome 8), the metabolic proto-oncogene and global glycolytic master regulator.

**Usage Notes**

To reproduce the Locus-Zoom grids (Supplemental Figure S1) or the Empirical Genomic Inventory (Table 1), run the Python plotting scripts within the same parent directory as these .tsv files. These scripts will be available with a Zenodo download associated with the manuscript.
