## Supplemental Dataset 2 for "State-Dependent Transcriptomic Collapse of the Brain’s Lactate and Ketone Thermodynamic Sensors in Schizophrenia": figure_2_panel_a_generator.py

```
import pandas as pd
import numpy as np
import matplotlib.pyplot as plt
import matplotlib as mpl
import seaborn as sns
import os

# ==========================================
# 0. Global Font Settings for SVG Export
# ==========================================
mpl.rcParams['svg.fonttype'] = 'none'
plt.rcParams['font.family'] = 'sans-serif'
plt.rcParams['font.sans-serif'] = ['Arial']
plt.rcParams['font.size'] = 10

def generate_panel_a():
    print("--- Generating Figure 2 Panel A: Baseline Forest Plot ---")
    
    # 1. Load the data
    input_file = "Figure2_Forest_Plot_Data.xlsx"
    try:
        df = pd.read_excel(input_file)
    except FileNotFoundError:
        print(f"⚠️ Error: Could not find '{input_file}'.")
        return
        
    # 2. Setup the visual theme (Nature/Cell style minimalism)
    sns.set_theme(style="whitegrid", context="paper")
    # Global font settings (Arial, editable SVG text) are defined at the top of the script
    
    # Sort the data so the most significant P-values are at the top of their blocks
    df = df.sort_values(by=['regulator', 'p-value'], ascending=[True, True])
    
    # 3. Create a figure with two stacked subplots (3' block and 5' block)
    fig, (ax1, ax2) = plt.subplots(2, 1, figsize=(10, 10), gridspec_kw={'height_ratios': [1, 1]}, sharex=True)
    
    # Assign stark, contrasting colors for the two receptors
    # Crimson for the Brake (HCAR1), Cobalt Blue for the Coolant (HCAR2)
    custom_palette = {"HCAR1": "#D62728", "HCAR2": "#1F77B4"} 
    
    # --- TOP PANEL: 3' Regulatory Skyscraper ---
    df_3p = df[df['regulator'] == "3'"]
    sns.stripplot(
        data=df_3p, 
        x="True_Disease_Delta", 
        y="rsID", 
        hue="gene",
        dodge=True,          # Offsets the dots slightly so they don't overlap
        orient="h", 
        alpha=0.85, 
        size=8, 
        palette=custom_palette, 
        ax=ax1
    )
    # Draw the central "Zero Effect" line
    ax1.axvline(x=0, color='black', linestyle='--', linewidth=1.5, zorder=0)
    ax1.set_title("3' Regulatory Skyscraper (Constitutive Lactate Brake Fracture)", fontsize=13, fontweight='bold', pad=10)
    ax1.set_xlabel("")
    ax1.set_ylabel("SCZ Risk Variant (rsID)", fontsize=11, fontweight='bold')
    
    # Clean up the legend and move it outside the plot
    ax1.legend(title="Target Gene", bbox_to_anchor=(1.02, 1), loc='upper left', frameon=True)

    # --- BOTTOM PANEL: 5' Regulatory Skyscraper ---
    df_5p = df[df['regulator'] == "5'"]
    sns.stripplot(
        data=df_5p, 
        x="True_Disease_Delta", 
        y="rsID", 
        hue="gene",
        dodge=True, 
        orient="h", 
        alpha=0.85, 
        size=8, 
        palette=custom_palette, 
        ax=ax2
    )
    # Draw the central "Zero Effect" line
    ax2.axvline(x=0, color='black', linestyle='--', linewidth=1.5, zorder=0)
    ax2.set_title("5' Regulatory Skyscraper (Dormant Baseline State)", fontsize=13, fontweight='bold', pad=10)
    ax2.set_xlabel("True Disease Delta (Δ Median INT Expression)", fontsize=12, fontweight='bold')
    ax2.set_ylabel("SCZ Risk Variant (rsID)", fontsize=11, fontweight='bold')
    
    # Remove redundant legend on bottom panel
    if ax2.get_legend():
        ax2.get_legend().remove()

    # 4. Final Formatting & Export
    plt.tight_layout()
    
    svg_out = "Figure_2_Panel_A_Forest_Plot.svg"
    png_out = "Figure_2_Panel_A_Forest_Plot.png"
    
    plt.savefig(svg_out, format="svg", bbox_inches="tight")
    plt.savefig(png_out, format="png", dpi=300, bbox_inches="tight")
    
    print(f"✅ Success! Figure generated.")
    print(f"📁 Vector SVG: {svg_out} (Use this for final manuscript/Illustrator)")
    print(f"📁 High-Res PNG: {png_out} (Use this for quick viewing/PPTs)")

if __name__ == "__main__":
    generate_panel_a()
```
