## Supplemental Dataset 2 for "State-Dependent Transcriptomic Collapse of the Brain’s Lactate and Ketone Thermodynamic Sensors in Schizophrenia": README.pdf

**Supplemental Dataset 2: Transcriptomic Fine-Mapping and Disease Directional Curation**

**Study:** State-Dependent Transcriptomic Collapse of the Brain's Lactate and Ketone Thermodynamic Sensors in Schizophrenia

**Author:** Bryan A. Krantz

**Overview**

This repository contains the data, manual curation records, and processing scripts required to bridge the SCZ structural variants (identified in Supplemental Dataset 1) to their functional transcriptomic consequences in the human cortex. This dataset provides the precise baseline eQTL data utilized to calculate the "True Disease Delta" and generate the transcriptomic double-dissociation forest plots (**Figure 2**).

**Methodology: Manual Curation and Directional Alignment**

Standard Genotype-Tissue Expression (GTEx V8) data portals and APIs frequently obscure nominal eQTL data and Inverse Normal Transformed (INT) expression medians if they do not meet highly conservative, genome-wide FDR thresholds. To perform rigorous, hypothesis-driven candidate testing on the *HCAR* tandem array, targeted manual curation was required.

1. **Manual Extraction:** The top SCZ risk variants from the 3' and 5' mutational skyscrapers were individually queried in the GTEx portal for the human cortex. Raw INT median expression values, exact genotype strings (e.g., CC, CT, TT), and sample sizes (n) were manually extracted from the UI violin plots to bypass automated masking.
2. **Disease Direction Alignment:** GTEx reports expression shifts strictly in terms of Reference vs. Alternate alleles. To determine the biophysical relevance to Schizophrenia, the raw GTEx delta was computationally cross-referenced against the specific SCZ risk allele from the PGC Wave 3 summary statistics. The resulting "True Disease Delta" isolates the exact transcriptomic up- or down-regulation driven specifically by the disease state.

**File Manifest**

**1. GWAS Input Targets (.tsv)**

These files contain the top index SNPs defining the 3' and 5' mutational skyscrapers, extracted directly from the PGC3 summary statistics (Dataset 1). They serve as the input queries for the GTEx portal.

- HCAR_3prime_cluster_top_SNPs.tsv: Top variants residing within the *HCAR1* (lactate brake) regulatory domain.
- HCAR_5prime_cluster_top_SNPs.tsv: Top variants residing within the *HCAR2* (BHB cooling switch) regulatory domain.

**2. Curated Transcriptomic Data (.xlsx)**

- cortex_GTEx_transcription_fine_data_medians_n_genotypes.xlsx: The master manual curation matrix. Contains the hand-extracted INT medians, genotypes, and cohort sizes (n) for all evaluated variants across the baseline human cortex.
- Figure2_Forest_Plot_Data.xlsx: The finalized, computed dataset combining the GWAS risk allele data with the GTEx expression data, resulting in the "True Disease Delta" utilized for plotting.

**3. Processing and Plotting Scripts (Python)**

- calculate_disease_direction.py: The computational logic script that imports the manual curation matrix, identifies the SCZ risk allele, and calculates the correct positive/negative shift (True Disease Delta) for the forest plot.
- figure_2_panel_a_generator.py: The visualization script utilizing matplotlib and seaborn to generate the dual-panel forest plot demonstrating the transcriptomic double dissociation.

**4. Output Figures**

- Figure_2_Panel_A_Forest_Plot.svg: Vector graphic output.
- Figure_2_Panel_A_Forest_Plot.png: High-resolution raster output.
