## Supplementary figures and images for "State-Dependent Transcriptomic Collapse of the Brain’s Lactate and Ketone Thermodynamic Sensors in Schizophrenia"

### Figure_2_Panel_A_Forest_Plot.png

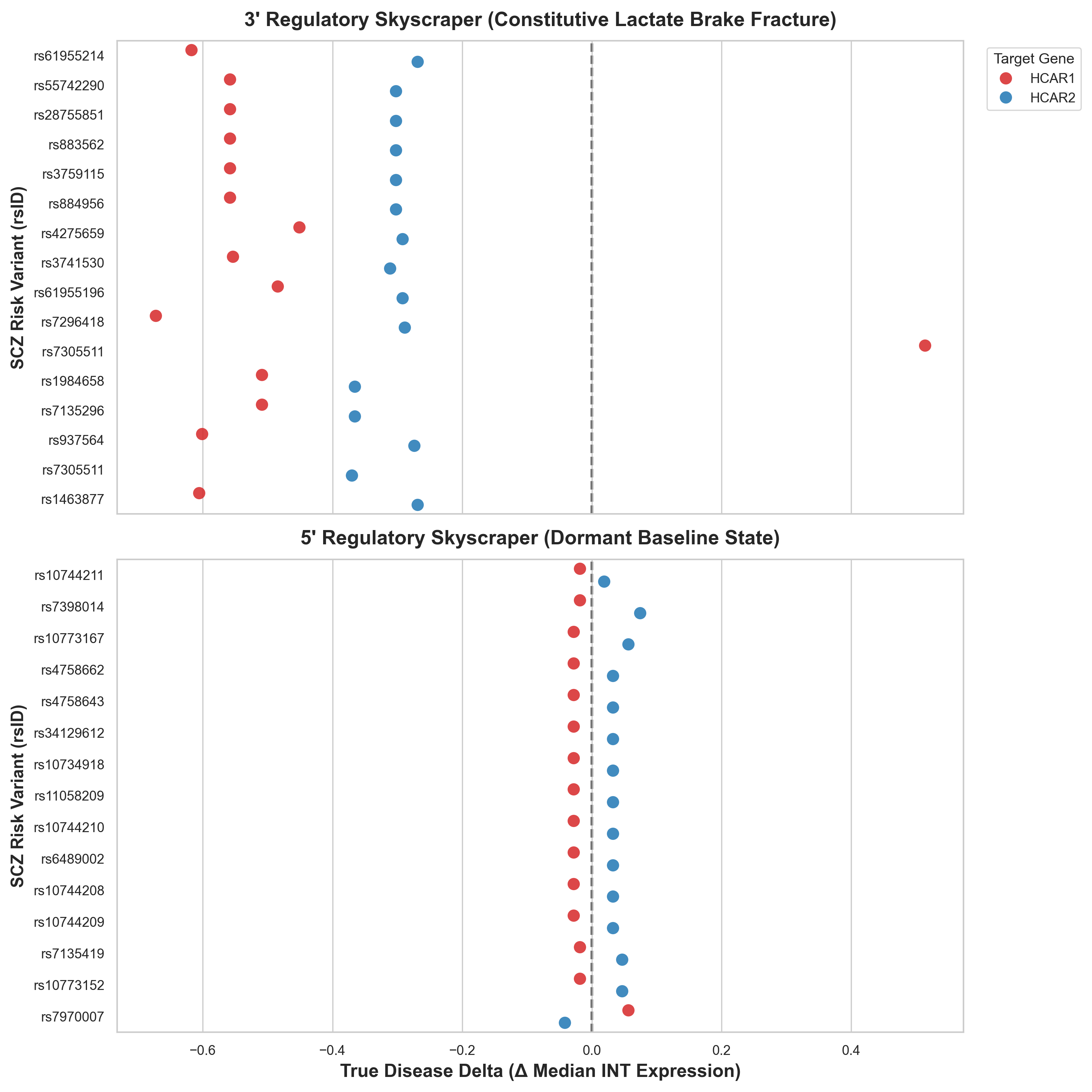

### Figure_3_Whole_Brain_Heatmap.png

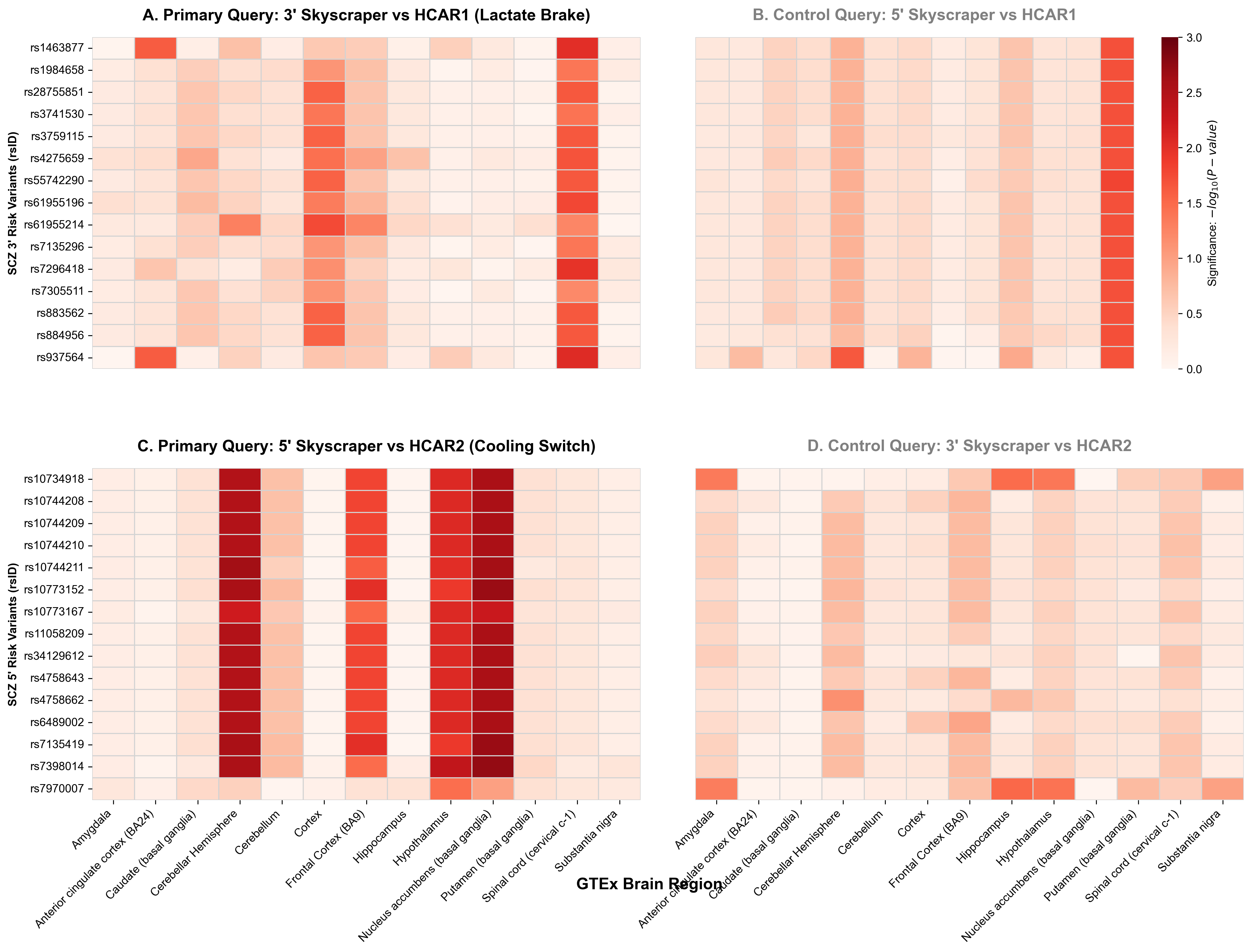

### Figure_S2_Brain_Dosage_Grid.png

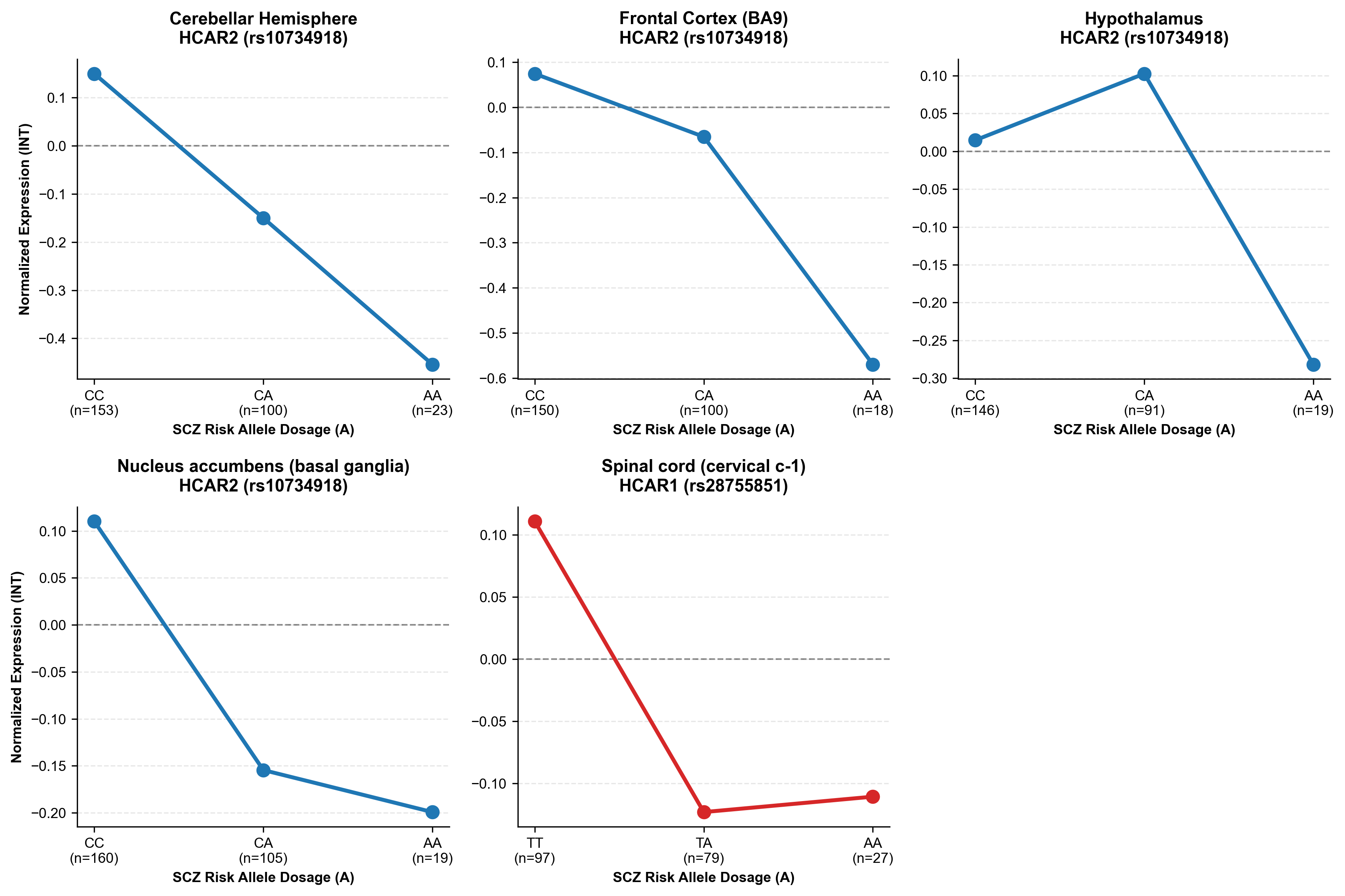
