## Supplemental Dataset 3 for "State-Dependent Transcriptomic Collapse of the Brain’s Lactate and Ketone Thermodynamic Sensors in Schizophrenia": figure_3_heatmap_generator.py

```
import pandas as pd
import numpy as np
import matplotlib.pyplot as plt
import seaborn as sns
import matplotlib as mpl
import os

# ==========================================
# 0. Global Font Settings for SVG Export
# ==========================================
mpl.rcParams['svg.fonttype'] = 'none'
plt.rcParams['font.family'] = 'sans-serif'
plt.rcParams['font.sans-serif'] = ['Arial']
plt.rcParams['font.size'] = 10

def generate_figure_3():
    print("--- Generating Figure 3: Whole-Brain Double Dissociation Heatmap ---")
    
    # Check for the correct filename (handles if you used the Aim1_ prefix or not)
    filename = "Aim1_Whole_Brain_Crown_Jewel_Matrix.xlsx"
    if not os.path.exists(filename):
        filename = "Whole_Brain_Crown_Jewel_Matrix.xlsx"
        if not os.path.exists(filename):
            print(f"⚠️ Error: Could not find '{filename}'.")
            return

    # Helper function to load, clean, and convert P-values to -log10(P)
    def prep_heatmap_data(sheet_name):
        df = pd.read_excel(filename, sheet_name=sheet_name, index_col=0)
        # Clean up the brain region names for the X-axis (Remove 'Brain - ' for clarity)
        df.columns = [str(col).replace('Brain - ', '') for col in df.columns]
        # Convert to -log10(P). Add a tiny constant to prevent log(0) errors.
        df_log = -np.log10(df.astype(float) + 1e-300)
        return df_log

    try:
        # Load the 4 matrices
        df_3p_hcar1 = prep_heatmap_data("Primary_3p_HCAR1")
        df_5p_hcar1 = prep_heatmap_data("Control_5p_HCAR1")
        df_5p_hcar2 = prep_heatmap_data("Primary_5p_HCAR2")
        df_3p_hcar2 = prep_heatmap_data("Control_3p_HCAR2")
    except ValueError as e:
        print(f"⚠️ Error reading sheets. Ensure all 4 Primary/Control tabs exist: {e}")
        return

    # Create the 2x2 Grid
    fig, axes = plt.subplots(2, 2, figsize=(18, 12), gridspec_kw={'wspace': 0.1, 'hspace': 0.3})
    
    # Establish a universal color scale (0 to 3.0). 
    # -log10(0.05) is ~1.3. So anything above 1.3 is significant (turns darker red).
    VMAX = 3.0
    VMIN = 0.0
    CMAP = "Reds"
    
    # --- Top Left: Primary HCAR1 (3' Skyscraper) ---
    sns.heatmap(df_3p_hcar1, ax=axes[0, 0], cmap=CMAP, vmin=VMIN, vmax=VMAX, cbar=False, 
                linewidths=0.5, linecolor='lightgray')
    axes[0, 0].set_title("A. Primary Query: 3' Skyscraper vs HCAR1 (Lactate Brake)", fontsize=14, fontweight='bold', pad=15)
    axes[0, 0].set_ylabel("SCZ 3' Risk Variants (rsID)", fontweight='bold')
    axes[0, 0].set_xticks([]) # Hide x-ticks for top row to keep it clean
    
    # --- Top Right: Control HCAR1 (5' Skyscraper) ---
    sns.heatmap(df_5p_hcar1, ax=axes[0, 1], cmap=CMAP, vmin=VMIN, vmax=VMAX, cbar=True,
                cbar_kws={'label': 'Significance: $-log_{10}(P-value)$'},
                linewidths=0.5, linecolor='lightgray')
    axes[0, 1].set_title("B. Control Query: 5' Skyscraper vs HCAR1", fontsize=14, fontweight='bold', pad=15, color='gray')
    axes[0, 1].set_ylabel("")
    axes[0, 1].set_xticks([])
    axes[0, 1].set_yticks([])

    # --- Bottom Left: Primary HCAR2 (5' Skyscraper) ---
    sns.heatmap(df_5p_hcar2, ax=axes[1, 0], cmap=CMAP, vmin=VMIN, vmax=VMAX, cbar=False,
                linewidths=0.5, linecolor='lightgray')
    axes[1, 0].set_title("C. Primary Query: 5' Skyscraper vs HCAR2 (Cooling Switch)", fontsize=14, fontweight='bold', pad=15)
    axes[1, 0].set_ylabel("SCZ 5' Risk Variants (rsID)", fontweight='bold')
    axes[1, 0].tick_params(axis='x', rotation=45)
    axes[1, 0].set_xticklabels(axes[1, 0].get_xticklabels(), ha='right')

    # --- Bottom Right: Control HCAR2 (3' Skyscraper) ---
    sns.heatmap(df_3p_hcar2, ax=axes[1, 1], cmap=CMAP, vmin=VMIN, vmax=VMAX, cbar=False,
                linewidths=0.5, linecolor='lightgray')
    axes[1, 1].set_title("D. Control Query: 3' Skyscraper vs HCAR2", fontsize=14, fontweight='bold', pad=15, color='gray')
    axes[1, 1].set_ylabel("")
    axes[1, 1].set_yticks([])
    axes[1, 1].tick_params(axis='x', rotation=45)
    axes[1, 1].set_xticklabels(axes[1, 1].get_xticklabels(), ha='right')

    # 4. Final Formatting & Export
    # Add a unified X-axis label
    fig.text(0.5, 0.02, 'GTEx Brain Region', ha='center', fontsize=14, fontweight='bold')
    
    svg_out = "Figure_3_Whole_Brain_Heatmap.svg"
    png_out = "Figure_3_Whole_Brain_Heatmap.png"
    
    plt.savefig(svg_out, format="svg", bbox_inches="tight")
    plt.savefig(png_out, format="png", dpi=300, bbox_inches="tight")
    
    print(f"✅ Success! Figure 3 generated.")
    print(f"📁 Vector SVG: {svg_out} (Fully editable in Inkscape)")
    print(f"📁 High-Res PNG: {png_out}")

if __name__ == "__main__":
    generate_figure_3()
```
