## Supplemental Dataset 3 for "State-Dependent Transcriptomic Collapse of the Brain’s Lactate and Ketone Thermodynamic Sensors in Schizophrenia": README.docx

**Supplemental Dataset 3: Whole-Brain Transcriptomic Footprint and Control eQTL Matrix**

**Study:** State-Dependent Transcriptomic Collapse of the Brain's Lactate and Ketone Thermodynamic Sensors in Schizophrenia

**Author:** Bryan A. Krantz

**Overview**

This repository contains the data, extraction logs, and processing scripts utilized to map the anatomical footprint of the *HCAR* tandem locus failure across the human central nervous system. By querying the 13 canonical brain regions in the GTEx V8 database, this dataset establishes the spatial specificity of the transcriptomic double-dissociation (e.g., 5' *HCAR2* fracture in the Nucleus Accumbens; 3' *HCAR1* fracture in the Cortex and Spinal Cord).

Crucially, this dataset also includes the exhaustive *trans*-eQTL control queries required to prove "Independent Flank Regulation," completely ruling out generic Linkage Disequilibrium (LD) artifacts.

**Methodology: GTEx Multi-Tissue Scraping and Parsing**

Because standard GTEx bulk downloads strictly mask nominal eQTLs failing to meet genome-wide FDR thresholds, high-resolution cross-tissue transcriptomic mapping required targeted manual queries.

1. **Batch Querying:** The index SCZ structural variants from the 3' and 5' mutational skyscrapers were systematically queried against both *HCAR1* and *HCAR2* across all 13 GTEx brain tissues.
2. **Raw Text Scraping:** The raw text outputs from the GTEx UI were compiled into master text logs to capture every nominal *P*-value across the neural circuits.
3. **Programmatic Parsing:** The custom whole_brain_master_scraper.py script utilized regex to parse these massive, unstructured text dumps, standardizing them into an interconnected genomic matrix (the "Crown Jewel" matrix) containing cleanly mapped rsID, Tissue, Gene, and P_Value columns.
4. **Heatmap Generation:** This matrix was then piped directly into seaborn plotting libraries to generate the 2x2 significance heatmap matrices (**Figure 3**).

**File Manifest**

**1. GTEx Query Inputs**

- GTEx_Whole_Brain_Queries.txt: The formatted text blocks used to query the primary hypotheses (3' Skyscraper vs. *HCAR1*; 5' Skyscraper vs. *HCAR2*) across the 13 brain regions.
- GTEx_Whole_Brain_Control_Queries.txt: The formatted text blocks used to query the converse *trans*-eQTL controls (3' Skyscraper vs. *HCAR2*; 5' Skyscraper vs. *HCAR1*).

**2. Raw Scrape Logs**

- entire_brain_control_GTEx_scrape.txt: The raw, unformatted text scrape containing the GTEx portal output for the control queries. (Note: The primary scrape text log is processed identically by the pipeline).

**3. Processing and Visualization Scripts (Python)**

- whole_brain_master_scraper.py: The master parsing script. It reads the raw GTEx text dumps, matches coordinates and alleles, and outputs a structured Excel workbook with dynamically generated Pivot Tables representing the complete whole-brain eQTL landscape.
- figure_3_heatmap_generator.py: The visualization script that reads the parsed matrix to render the 2x2 multi-panel heatmap shown in Figure 3.

**4. Processed Data Matrices**

- Whole_Brain_Crown_Jewel.tsv: The flat, tabular output of all parsed eQTLs across the 13 brain regions.
- Whole_Brain_Crown_Jewel_Matrix.xlsx: The finalized computational matrix. Contains the flat data alongside four discrete pivot tables:
  - Primary_3p_HCAR1
  - Primary_5p_HCAR2
  - Control_5p_HCAR1 (Proving Independent Regulation)
  - Control_3p_HCAR2 (Proving Independent Regulation)

**5. Output Figures**

- Figure_3_Whole_Brain_Heatmap.svg: Vector graphic output (2x2 Panel).
- Figure_3_Whole_Brain_Heatmap.png: High-resolution raster output (2x2 Panel).
