## Supplemental Dataset 3 for "State-Dependent Transcriptomic Collapse of the Brain’s Lactate and Ketone Thermodynamic Sensors in Schizophrenia": whole_brain_master_scraper.py

```
import re
import os
import pandas as pd

# The two files you scraped
FILE_PRIMARY = "entire_brain_GTEx_scrape.txt"
FILE_CONTROL = "entire_brain_control_GTEx_scrape.txt"

standard_columns = [
    "Regulatory_Region", 
    "Gene", 
    "Ensembl_ID", 
    "rsID", 
    "Coordinates_b38", 
    "Tissue", 
    "P_Value"
]

# Canonical 13 GTEx Brain Regions (Strictly Alphabetical)
CANONICAL_BRAIN_TISSUES = [
    "Brain - Amygdala",
    "Brain - Anterior cingulate cortex (BA24)",
    "Brain - Caudate (basal ganglia)",
    "Brain - Cerebellar Hemisphere",
    "Brain - Cerebellum",
    "Brain - Cortex",
    "Brain - Frontal Cortex (BA9)",
    "Brain - Hippocampus",
    "Brain - Hypothalamus",
    "Brain - Nucleus accumbens (basal ganglia)",
    "Brain - Putamen (basal ganglia)",
    "Brain - Spinal cord (cervical c-1)",
    "Brain - Substantia nigra"
]

def parse_massive_brain_dump(filepath, is_control_file=False):
    if not os.path.exists(filepath):
        print(f"⚠️ Warning: Could not find '{filepath}'. Skipping.")
        return pd.DataFrame()

    with open(filepath, 'r', encoding='utf-8') as f:
        raw_text = f.read()

    blocks = re.split(r'\n(?=HCAR[12]\s+\()', '\n' + raw_text.strip())
    extracted_data = []
    
    for block in blocks:
        if not block.strip() or "pvalue" not in block:
            continue
            
        gene_match = re.search(r'(HCAR[12])', block)
        ensembl_match = re.search(r'(ENSG[\d\.]+)', block)
        rsid_match = re.search(r'(rs\d+)', block)
        coord_match = re.search(r'(chr\d+_\d+_[A-Z]_[A-Z]_b38)', block)
        pvals = re.findall(r'pvalue:\s+([\d\.e\-]+)', block)
        
        if gene_match and rsid_match and pvals:
            gene = gene_match.group(1)
            rsid = rsid_match.group(1)
            ensembl = ensembl_match.group(1) if ensembl_match else "Unknown"
            coord = coord_match.group(1) if coord_match else "Unknown"
            
            # --- Dynamic Regulatory Assignment ---
            if not is_control_file:
                # Primary Query: HCAR1 = 3', HCAR2 = 5'
                region = "3' Regulatory Skyscraper" if gene == 'HCAR1' else "5' Regulatory Skyscraper"
            else:
                # Control Query (The Sanity Check): HCAR1 = 5', HCAR2 = 3'
                region = "5' Regulatory Skyscraper" if gene == 'HCAR1' else "3' Regulatory Skyscraper"
            
            if len(pvals) == len(CANONICAL_BRAIN_TISSUES):
                for i, pval_str in enumerate(pvals):
                    extracted_data.append({
                        "Regulatory_Region": region,
                        "Gene": gene,
                        "Ensembl_ID": ensembl,
                        "rsID": rsid,
                        "Coordinates_b38": coord,
                        "Tissue": CANONICAL_BRAIN_TISSUES[i],
                        "P_Value": float(pval_str)
                    })
            else:
                print(f"⚠️ Note: rsID {rsid} ({gene}) in {filepath} yielded {len(pvals)} P-values (Likely a GTEx rare-genotype drop).")
                for i, pval_str in enumerate(pvals):
                    extracted_data.append({
                        "Regulatory_Region": region,
                        "Gene": gene,
                        "Ensembl_ID": ensembl,
                        "rsID": rsid,
                        "Coordinates_b38": coord,
                        "Tissue": f"Unknown Alignment Region {i+1}",
                        "P_Value": float(pval_str)
                    })

    df = pd.DataFrame(extracted_data)
    if not df.empty:
        df = df[standard_columns]
    return df

def main():
    print("--- Assembling the Whole-Brain 'Crown Jewel' Database ---")
    
    # 1. Parse both the primary and control files
    df_primary = parse_massive_brain_dump(FILE_PRIMARY, is_control_file=False)
    df_control = parse_massive_brain_dump(FILE_CONTROL, is_control_file=True)
    
    # 2. Combine into a single master flat database
    master_df = pd.concat([df_primary, df_control], ignore_index=True)
    
    if not master_df.empty:
        master_df = master_df.sort_values(by=["Regulatory_Region", "Gene", "Tissue", "P_Value"])
        
        # Output 1: Master TSV
        tsv_out = "Whole_Brain_Crown_Jewel.tsv"
        master_df.to_csv(tsv_out, sep='\t', index=False)
        print(f"📁 Created Master TSV Database: {tsv_out}")
        
        # Output 2: The 5-Tab Excel Matrix
        excel_out = "Whole_Brain_Crown_Jewel_Matrix.xlsx"
        try:
            with pd.ExcelWriter(excel_out, engine='openpyxl') as writer:
                # Primary Tabs
                df_3p_hcar1 = master_df[(master_df['Regulatory_Region'] == "3' Regulatory Skyscraper") & (master_df['Gene'] == "HCAR1")]
                if not df_3p_hcar1.empty:
                    df_3p_hcar1.pivot_table(index="rsID", columns="Tissue", values="P_Value").to_excel(writer, sheet_name="Primary_3p_HCAR1")
                
                df_5p_hcar2 = master_df[(master_df['Regulatory_Region'] == "5' Regulatory Skyscraper") & (master_df['Gene'] == "HCAR2")]
                if not df_5p_hcar2.empty:
                    df_5p_hcar2.pivot_table(index="rsID", columns="Tissue", values="P_Value").to_excel(writer, sheet_name="Primary_5p_HCAR2")
                
                # Control Tabs
                df_5p_hcar1 = master_df[(master_df['Regulatory_Region'] == "5' Regulatory Skyscraper") & (master_df['Gene'] == "HCAR1")]
                if not df_5p_hcar1.empty:
                    df_5p_hcar1.pivot_table(index="rsID", columns="Tissue", values="P_Value").to_excel(writer, sheet_name="Control_5p_HCAR1")
                    
                df_3p_hcar2 = master_df[(master_df['Regulatory_Region'] == "3' Regulatory Skyscraper") & (master_df['Gene'] == "HCAR2")]
                if not df_3p_hcar2.empty:
                    df_3p_hcar2.pivot_table(index="rsID", columns="Tissue", values="P_Value").to_excel(writer, sheet_name="Control_3p_HCAR2")

                # Raw Data
                master_df.to_excel(writer, sheet_name="Raw_Flat_Data", index=False)
                
            print(f"📊 Created 'Crown Jewel' Pivot Matrix: {excel_out}")
            print(f"✅ Extracted a total of {len(master_df)} neural variants across primary and control targets.")
        except ModuleNotFoundError:
            print("\n⚠️ Note: Install 'openpyxl' (pip install openpyxl) to generate the Excel matrices.")
            
    else:
        print("❌ No data parsed. Check that your .txt files are in the same folder.")

if __name__ == "__main__":
    main()
```
