## Supplemental Dataset 4 for "State-Dependent Transcriptomic Collapse of the Brain’s Lactate and Ketone Thermodynamic Sensors in Schizophrenia": phase4_systemic_control_query.py

```
import csv
import os

# We will use the same 5' and 3' top SNPs extracted during Phase 1
input_5prime = "HCAR_5prime_cluster_top_SNPs.tsv"
input_3prime = "HCAR_3prime_cluster_top_SNPs.tsv"

# Ensembl IDs
HCAR1_VERSION = "ENSG00000196917.6"  # HCAR1 (Lactate Brake)
HCAR2_VERSION = "ENSG00000182782.8"  # HCAR2 (BHB Cooling Switch - Niacin Receptor)

def extract_rsids(filepath):
    rsids = []
    if os.path.exists(filepath):
        with open(filepath, 'r') as f:
            reader = csv.DictReader(f, delimiter='\t')
            for row in reader:
                if row['ID'].startswith('rs'):
                    rsids.append(row['ID'])
    return rsids[:15] # GTEx UI limits to chunks, 15 is safe

def main():
    print("Generating GTEx UI Queries for Systemic Control Tissues...")
    
    rsids_5p = extract_rsids(input_5prime)
    rsids_3p = extract_rsids(input_3prime)
    
    if not rsids_5p or not rsids_3p:
        print("Error: Could not find Phase 1 TSV files. Make sure they are in the directory.")
        return

    output_file = "GTEx_Systemic_Control_Queries.txt"
    
    with open(output_file, 'w') as f:
        f.write("--- SYSTEMIC CONTROL QUERIES (The 'Missing Niacin Flush' Test) ---\n")
        f.write("Hypothesis: The 5' and 3' structural variants systemically downregulate HCAR2 in the skin and vasculature, physically explaining the historic absence of the Niacin Flush in SCZ patients.\n\n")
        
        f.write("TARGET TISSUES TO SELECT IN GTEx UI:\n")
        f.write("- Skin - Sun Exposed (Lower leg)\n")
        f.write("- Skin - Not Sun Exposed (Suprapubic)\n")
        f.write("- Whole Blood\n")
        f.write("- Cells - Cultured fibroblasts\n\n")
        f.write("=====================================================\n\n")
        
        f.write("BLOCK 1: 5' Skyscraper vs HCAR2 (Niacin Receptor)\n")
        f.write("Tests if the 5' variant breaks HCAR2 systemically.\n")
        for rsid in rsids_5p:
            f.write(f"{HCAR2_VERSION},{rsid}\n")
            
        f.write("\n=====================================================\n\n")
        
        f.write("BLOCK 2: 3' Skyscraper vs HCAR1 (Systemic Lactate Brake)\n")
        f.write("Tests if the 3' variant breaks HCAR1 systemically.\n")
        for rsid in rsids_3p:
            f.write(f"{HCAR1_VERSION},{rsid}\n")

    print(f"Exported queries to {output_file}.")
    print("Instructions: Paste these into the GTEx eQTL Dashboard and select Skin/Blood as the target tissues.")

if __name__ == "__main__":
    main()
```
