## Supplemental Dataset 4 for "State-Dependent Transcriptomic Collapse of the Brain’s Lactate and Ketone Thermodynamic Sensors in Schizophrenia": README.docx

**Supplemental Dataset 4: Systemic Transcriptomic Footprint and Peripheral Pleiotropy eQTL Matrix**

**Study:** State-Dependent Transcriptomic Collapse of the Brain's Lactate and Ketone Thermodynamic Sensors in Schizophrenia

**Author:** Bryan A. Krantz

**Overview**

This repository contains the raw data, extraction logs, and processing scripts utilized to map the systemic and peripheral footprint of the *HCAR* tandem locus failure across highly metabolic human tissues. By querying specific non-brain tissues in the GTEx V8 database, this dataset establishes the robust peripheral consequences of the thermodynamic fracture.

Critically, this data maps the systemic lipid dysregulation associated with the 5' *HCAR2* fracture (in adipose and fibroblast tissues) and confirms the presence of "Antagonistic Pleiotropy" regarding the 3' *HCAR1* fracture—demonstrating a massive, dose-dependent upregulation of the lactate shuttle in the testis that likely provides the reproductive advantage maintaining this architecture in the human gene pool.

**Methodology: GTEx Peripheral Tissue Scraping and Parsing**

Because standard GTEx bulk downloads strictly mask nominal eQTLs failing to meet genome-wide FDR thresholds, this systemic transcriptomic mapping required targeted manual queries across specifically chosen peripheral control tissues.

1. **Target Identification:** The index SCZ structural variants from the 3' and 5' mutational skyscrapers were queried against *HCAR1* and *HCAR2* across highly metabolic tissues (e.g., Testis, Adipose, Liver, Breast, Fibroblasts) and distinct control tissues (e.g., Ovary, Whole Blood, Skin).
2. **Raw Text Scraping:** The raw text outputs from the GTEx UI were compiled into individual text logs for each tissue and target combination.
3. **Programmatic Parsing:** The custom systemic_control_master_scraper.py script was utilized to iterate through the individual text files. Using regex, it parsed the unstructured text dumps and standardized them into a centralized, cross-tissue genomic matrix mapping the exact rsID, Tissue, Regulatory_Region, and P_Value.
4. **Matrix Consolidation:** The parser automatically sorts the data by significance and exports both a flat TSV file and a highly structured, multi-tab Excel workbook for rapid analysis and figure generation (**Figure 4**).

**File Manifest**

**1. GWAS Inputs & Query Generation**

- HCAR_3prime_cluster_top_SNPs.tsv: The top index SNPs from the 3' regulatory skyscraper (PGC Wave 3).
- HCAR_5prime_cluster_top_SNPs.tsv: The top index SNPs from the 5' mutational cluster (PGC Wave 3).
- phase4_systemic_control_query.py: Script utilized to batch-generate the precise query strings for the GTEx portal.
- GTEx_Systemic_Control_Queries.txt: The formatted text blocks used for copy-pasting queries into the GTEx interface.

**2. Raw Scrape Logs (Tissue-Specific)**

Individual text scrapes containing the GTEx portal output, divided by target tissue and regulatory region:

- adipose_subcutaneous_3prime_HCAR1_GTEx_scrape.txt & adipose_subcutaneous_5prime_HCAR2_GTEx_scrape.txt
- adipose_visceral_3prime_HCAR1_GTEx_scrape.txt & adipose_visceral_5prime_HCAR2_GTEx_scrape.txt
- breast_3prime_HCAR1_GTEx_scrape.txt & breast_5prime_HCAR2_GTEx_scrape.txt
- fibroblasts_3prime_HCAR1_GTEx_scrape.txt & fibroblasts_5prime_HCAR2_GTEx_scrape.txt
- liver_3prime_HCAR1_GTEx_scrape.txt & liver_5prime_HCAR2_GTEx_scrape.txt
- ovary_3prime_HCAR1_GTEx_scrape.txt & ovary_5prime_HCAR2_GTEx_scrape.txt
- skin_lower_leg_sun_exposed_... & skin_suprapubic_not_sun_exposed_... (Control scrapes)
- testis_3prime_HCAR1_GTEx_scrape.txt & testis_5prime_HCAR2_GTEx_scrape.txt (Pleiotropy targets)
- whole_blood_3prime_HCAR1_GTEx_scrape.txt & whole_blood_5prime_HCAR2_GTEx_scrape.txt

**3. Processing Scripts (Python)**

- systemic_control_master_scraper.py: The master parsing script. It maps the 14 individual text files to their corresponding metadata, extracts the coordinate/significance data, and outputs the consolidated master matrices.

**4. Processed Data Matrices**

- Systemic_Controls_Master.tsv: The flat, tabular output of all parsed eQTLs across the peripheral tissues, sorted by target region and significance.
- Systemic_Controls_Master.xlsx: The finalized computational matrix organized into distinct tabs (e.g., Testis_3p_HCAR1, Adipose_5p_HCAR2) for immediate visualization and downstream plotting.
