## Supplemental Dataset 4 for "State-Dependent Transcriptomic Collapse of the Brain’s Lactate and Ketone Thermodynamic Sensors in Schizophrenia": systemic_control_master_scraper.py

```
import os
import re
import pandas as pd

# The 14 files and their corresponding metadata. 
# Tab names are kept under 31 characters to comply with Excel limits.
file_mapping = {
    "skin_lower_leg_sun_exposed_5prime_HCAR2_GTEx_scrape.txt": {
        "Regulatory_Region": "5' Regulatory Skyscraper", "Tissue": "Skin - Sun Exposed (Lower Leg)", "Tab_Name": "SkinSun_5p_HCAR2"
    },
    "skin_lower_leg_sun_exposed_3prime_HCAR1_GTEx_scrape.txt": {
        "Regulatory_Region": "3' Regulatory Skyscraper", "Tissue": "Skin - Sun Exposed (Lower Leg)", "Tab_Name": "SkinSun_3p_HCAR1"
    },
    "skin_suprapubic_not_sun_exposed_5prime_HCAR2_GTEx_scrape.txt": {
        "Regulatory_Region": "5' Regulatory Skyscraper", "Tissue": "Skin - Unexposed (Suprapubic)", "Tab_Name": "SkinUnexp_5p_HCAR2"
    },
    "skin_suprapubic_not_sun_exposed_3prime_HCAR1_GTEx_scrape.txt": {
        "Regulatory_Region": "3' Regulatory Skyscraper", "Tissue": "Skin - Unexposed (Suprapubic)", "Tab_Name": "SkinUnexp_3p_HCAR1"
    },
    "whole_blood_5prime_HCAR2_GTEx_scrape.txt": {
        "Regulatory_Region": "5' Regulatory Skyscraper", "Tissue": "Whole Blood", "Tab_Name": "Blood_5p_HCAR2"
    },
    "whole_blood_3prime_HCAR1_GTEx_scrape.txt": {
        "Regulatory_Region": "3' Regulatory Skyscraper", "Tissue": "Whole Blood", "Tab_Name": "Blood_3p_HCAR1"
    },
    "fibroblasts_5prime_HCAR2_GTEx_scrape.txt": {
        "Regulatory_Region": "5' Regulatory Skyscraper", "Tissue": "Cultured Fibroblasts", "Tab_Name": "Fibroblast_5p_HCAR2"
    },
    "fibroblasts_3prime_HCAR1_GTEx_scrape.txt": {
        "Regulatory_Region": "3' Regulatory Skyscraper", "Tissue": "Cultured Fibroblasts", "Tab_Name": "Fibroblast_3p_HCAR1"
    },
    "adipose_subcutaneous_5prime_HCAR2_GTEx_scrape.txt": {
        "Regulatory_Region": "5' Regulatory Skyscraper", "Tissue": "Adipose - Subcutaneous", "Tab_Name": "AdiposeSub_5p_HCAR2"
    },
    "adipose_subcutaneous_3prime_HCAR1_GTEx_scrape.txt": {
        "Regulatory_Region": "3' Regulatory Skyscraper", "Tissue": "Adipose - Subcutaneous", "Tab_Name": "AdiposeSub_3p_HCAR1"
    },
    "adipose_visceral_5prime_HCAR2_GTEx_scrape.txt": {
        "Regulatory_Region": "5' Regulatory Skyscraper", "Tissue": "Adipose - Visceral (Omentum)", "Tab_Name": "AdiposeVis_5p_HCAR2"
    },
    "adipose_visceral_3prime_HCAR1_GTEx_scrape.txt": {
        "Regulatory_Region": "3' Regulatory Skyscraper", "Tissue": "Adipose - Visceral (Omentum)", "Tab_Name": "AdiposeVis_3p_HCAR1"
    },
    "liver_5prime_HCAR2_GTEx_scrape.txt": {
        "Regulatory_Region": "5' Regulatory Skyscraper", "Tissue": "Liver", "Tab_Name": "Liver_5p_HCAR2"
    },
    "liver_3prime_HCAR1_GTEx_scrape.txt": {
        "Regulatory_Region": "3' Regulatory Skyscraper", "Tissue": "Liver", "Tab_Name": "Liver_3p_HCAR1"
    },
    "breast_5prime_HCAR2_GTEx_scrape.txt": {
        "Regulatory_Region": "5' Regulatory Skyscraper", "Tissue": "Breast - Mammary Tissue", "Tab_Name": "Breast_5p_HCAR2"
    },
    "breast_3prime_HCAR1_GTEx_scrape.txt": {
        "Regulatory_Region": "3' Regulatory Skyscraper", "Tissue": "Breast - Mammary Tissue", "Tab_Name": "Breast_3p_HCAR1"
    },
    "ovary_5prime_HCAR2_GTEx_scrape.txt": {
        "Regulatory_Region": "5' Regulatory Skyscraper", "Tissue": "Ovary", "Tab_Name": "Ovary_5p_HCAR2"
    },
    "ovary_3prime_HCAR1_GTEx_scrape.txt": {
        "Regulatory_Region": "3' Regulatory Skyscraper", "Tissue": "Ovary", "Tab_Name": "Ovary_3p_HCAR1"
    },    
    "testis_5prime_HCAR2_GTEx_scrape.txt": {
        "Regulatory_Region": "5' Regulatory Skyscraper", "Tissue": "Testis", "Tab_Name": "Testis_5p_HCAR2"
    },
    "testis_3prime_HCAR1_GTEx_scrape.txt": {
        "Regulatory_Region": "3' Regulatory Skyscraper", "Tissue": "Testis", "Tab_Name": "Testis_3p_HCAR1"
    },  
}

standard_columns = [
    "Regulatory_Region", 
    "Gene", 
    "Ensembl_ID", 
    "rsID", 
    "Coordinates_b38", 
    "Tissue", 
    "P_Value"
]

def process_systemic_scrapes():
    # Regex designed to jump directly to the payload, ignoring weird headers or footers
    # Group 1: Gene, Group 2: Ensembl, Group 3: rsID, Group 4: b38 Coords, Group 5: P-value
    extraction_pattern = re.compile(
        r'(HCAR[12])\s+\((ENSG[\d\.]+)\)\s+and\s+(rs\d+)\s+\((chr\d+_\d+_[A-Z]_[A-Z]_b38)\)\s*pvalue:\s+([\d\.e\-]+)'
    )
    
    all_dataframes = {}
    master_records = []
    
    print("--- Starting Systemic Tissue Scrape Compilation ---")
    
    for filename, meta in file_mapping.items():
        if not os.path.exists(filename):
            print(f"⚠️ Warning: File '{filename}' not found. Skipping.")
            continue
            
        with open(filename, 'r', encoding='utf-8') as f:
            content = f.read()
            
        matches = extraction_pattern.findall(content)
        
        # Handle the "Stubs" gracefully
        if not matches:
            print(f"⏩ Skipped (Stub/Empty): {filename}")
            continue
            
        print(f"✅ Parsed {len(matches)} variants from: {filename}")
        
        file_data = []
        for match in matches:
            file_data.append({
                "Regulatory_Region": meta["Regulatory_Region"],
                "Gene": match[0],
                "Ensembl_ID": match[1],
                "rsID": match[2],
                "Coordinates_b38": match[3],
                "Tissue": meta["Tissue"],
                "P_Value": float(match[4])
            })
            
        # Convert to DataFrame, ensure standardized column order, and sort by significance
        df = pd.DataFrame(file_data)[standard_columns]
        df = df.sort_values(by="P_Value", ascending=True)
        
        all_dataframes[meta["Tab_Name"]] = df
        master_records.append(df)
        
    # --- Output Generation ---
    if master_records:
        master_df = pd.concat(master_records, ignore_index=True)
        master_df = master_df.sort_values(by=["Regulatory_Region", "Gene", "Tissue", "P_Value"])
        
        # 1. Master TSV
        tsv_filename = "Systemic_Controls_Master.tsv"
        master_df.to_csv(tsv_filename, sep='\t', index=False)
        print(f"\n📁 Master TSV created: {tsv_filename}")
        
        # 2. Multi-Tab Excel
        excel_filename = "Systemic_Controls_Master.xlsx"
        try:
            with pd.ExcelWriter(excel_filename, engine='openpyxl') as writer:
                for tab_name, df in all_dataframes.items():
                    df.to_excel(writer, sheet_name=tab_name, index=False)
            print(f"📊 Multi-Tab Excel created: {excel_filename}")
        except ModuleNotFoundError:
            print("\n⚠️ Note: Install 'openpyxl' (pip install openpyxl) to generate the Excel file.")
            
    else:
        print("\n❌ No data was extracted from any file. Check file contents.")

if __name__ == "__main__":
    process_systemic_scrapes()
```
