## Supplemental Dataset 5 for "State-Dependent Transcriptomic Collapse of the Brain’s Lactate and Ketone Thermodynamic Sensors in Schizophrenia": calculate_subcortical_disease_direction.py

```
import pandas as pd
import os

# Updated to use the new combined file
MANUAL_EXTRACT_FILE = "cortex&other_brain_areas_GTEx_transcription_fine_data_medians_n_genotypes.xlsx"
SHEET_NAME = "other_brain_areas"

# The original GWAS data to cross-reference the SCZ Risk Allele
GWAS_3PRIME = "HCAR_3prime_cluster_top_SNPs.tsv"
GWAS_5PRIME = "HCAR_5prime_cluster_top_SNPs.tsv"

def calculate_subcortical_disease_effect():
    print("--- Calculating True Biological Direction of Transcription for Subcortical Areas ---")
    
    if not os.path.exists(MANUAL_EXTRACT_FILE):
        print(f"⚠️ Error: Please ensure '{MANUAL_EXTRACT_FILE}' is in the directory.")
        return

    # 1. Load the User's manual extractions specifically from the target sheet
    try:
        df_extracted = pd.read_excel(MANUAL_EXTRACT_FILE, sheet_name=SHEET_NAME)
    except Exception as e:
        print(f"⚠️ Error reading sheet '{SHEET_NAME}': {e}")
        return
    
    # 2. Load the GWAS data to get the A1 (Risk) and A2 (Non-Risk) alleles
    try:
        df_3p = pd.read_csv(GWAS_3PRIME, sep='\t')
        df_5p = pd.read_csv(GWAS_5PRIME, sep='\t')
        df_gwas = pd.concat([df_3p, df_5p])[['ID', 'A1', 'A2']]
        df_gwas = df_gwas.rename(columns={'ID': 'rsID', 'A1': 'SCZ_Risk_Allele', 'A2': 'Safe_Allele'})
    except FileNotFoundError:
        print(f"⚠️ Error: Could not find GWAS TSV files. Make sure they are in the directory.")
        return

    # 3. Merge the datasets
    df_merged = pd.merge(df_extracted, df_gwas, on='rsID', how='left')

    # 4. Calculate the True Disease Delta
    true_deltas = []
    directions = []
    
    for idx, row in df_merged.iterrows():
        # The raw GTEx Delta (Right Violin - Left Violin)
        raw_delta = row['median_3'] - row['median_1']
        
        # GTEx always plots Ref -> Alt. 
        # We extract the single nucleotide from the homozygous strings (e.g., 'GG' -> 'G')
        ref_allele = str(row['genotype_1'])[0]
        alt_allele = str(row['genotype_3'])[0]
        
        if row['SCZ_Risk_Allele'] == alt_allele:
            # The GTEx plot matches the disease direction.
            true_delta = raw_delta
        elif row['SCZ_Risk_Allele'] == ref_allele:
            # The GTEx plot is backwards relative to the disease. We flip the sign.
            true_delta = -raw_delta
        else:
            # Fallback if alleles are ambiguous (e.g., strand flips)
            true_delta = raw_delta
            
        true_deltas.append(true_delta)
        
        # Human-readable direction
        if true_delta < -0.1:
            directions.append("DOWN-regulated in SCZ")
        elif true_delta > 0.1:
            directions.append("UP-regulated in SCZ")
        else:
            directions.append("No Effect (Flat)")

    df_merged['True_Disease_Delta'] = true_deltas
    df_merged['Biological_Consequence'] = directions

    # 5. Export to a NEW file so it doesn't overwrite Figure 2 data
    output_file = "Subcortical_Disease_Direction_Calculated.xlsx"
    df_merged.to_excel(output_file, index=False)
    
    print(f"✅ Successfully calculated the true biological direction for {len(df_merged)} subcortical variants!")
    print(f"📁 Data saved to: {output_file}")

if __name__ == "__main__":
    calculate_subcortical_disease_effect()
```
