## Supplemental Dataset 5 for "State-Dependent Transcriptomic Collapse of the Brain’s Lactate and Ketone Thermodynamic Sensors in Schizophrenia": figure_s2_subcortical_dosage.py

```
import pandas as pd
import matplotlib.pyplot as plt
import matplotlib as mpl
import os

# ==========================================
# 0. Global Font Settings for SVG Export
# ==========================================
mpl.rcParams['svg.fonttype'] = 'none'
plt.rcParams['font.family'] = 'sans-serif'
plt.rcParams['font.sans-serif'] = ['Arial']
plt.rcParams['font.size'] = 9

def generate_figure_s2():
    print("--- Generating Figure S2: Subcortical & Regional Dosage Grids ---")
    
    file_name = "Subcortical_Disease_Direction_Calculated.xlsx"
    
    if not os.path.exists(file_name):
        print(f"⚠️ Error: Could not find '{file_name}'.")
        return
        
    try:
        # Default pandas export uses 'Sheet1', so we drop the specific sheet_name argument
        df = pd.read_excel(file_name)
    except Exception as e:
        print(f"⚠️ Error reading file: {e}")
        return

    # Create a 2x3 grid
    fig, axes = plt.subplots(2, 3, figsize=(12, 8))
    axes = axes.flatten()
    
    # Custom colors to match our theme (Red for HCAR1, Blue for HCAR2)
    color_map = {'HCAR1': '#D62728', 'HCAR2': '#1F77B4'}
    
    # We will plot up to 6 regions (2x3 grid)
    plot_count = min(len(df), 6)
    
    for i in range(plot_count):
        row = df.iloc[i]
        ax = axes[i]
        
        gene = str(row.get('gene', 'Unknown'))
        tissue = str(row.get('tissue', 'Unknown Brain Region'))
        rsid = str(row.get('rsID', 'Unknown rsID'))
        risk_allele = str(row.get('SCZ_Risk_Allele', '')).strip()
        
        # GTEx plots Ref/Ref, Ref/Alt, Alt/Alt
        ref_allele = str(row.get('genotype_1', 'XX'))[0]
        alt_allele = str(row.get('genotype_3', 'YY'))[0]
        
        # Orient the dosage based on the SCZ Risk Allele
        # If the risk allele is the Alternate allele (normal GTEx orientation)
        if risk_allele == alt_allele:
            y_vals = [row['median_1'], row['median_2'], row['median_3']]
            n_vals = [row['n_1'], row['n_2'], row['n_3']]
            x_labels = [f"{ref_allele}{ref_allele}\n(n={n_vals[0]})", 
                        f"{ref_allele}{alt_allele}\n(n={n_vals[1]})", 
                        f"{alt_allele}{alt_allele}\n(n={n_vals[2]})"]
        # If the risk allele is the Reference allele (we must flip the plot)
        elif risk_allele == ref_allele:
            y_vals = [row['median_3'], row['median_2'], row['median_1']]
            n_vals = [row['n_3'], row['n_2'], row['n_1']]
            x_labels = [f"{alt_allele}{alt_allele}\n(n={n_vals[0]})", 
                        f"{alt_allele}{ref_allele}\n(n={n_vals[1]})", 
                        f"{ref_allele}{ref_allele}\n(n={n_vals[2]})"]
        else:
            print(f"⚠️ Warning: Risk allele '{risk_allele}' doesn't match {ref_allele} or {alt_allele} for {rsid}. Skipping panel.")
            continue
            
        color = color_map.get(gene, '#333333')
        
        # Plot the line
        ax.plot([0, 1, 2], y_vals, marker='o', markersize=8, color=color, linewidth=2.5, zorder=3)
        ax.axhline(y=0, color='gray', linestyle='--', linewidth=1, zorder=1)
        
        # Formatting
        ax.set_title(f"{tissue}\n{gene} ({rsid})", fontsize=11, fontweight='bold', pad=10)
        ax.set_xticks([0, 1, 2])
        ax.set_xticklabels(x_labels)
        
        if i >= 0:
            ax.set_xlabel(f"SCZ Risk Allele Dosage ({risk_allele})", fontweight='bold')
        if i % 3 == 0:
            ax.set_ylabel("Normalized Expression (INT)", fontweight='bold')
            
        ax.spines['top'].set_visible(False)
        ax.spines['right'].set_visible(False)
        ax.grid(axis='y', linestyle='--', alpha=0.3)

    # Hide any unused subplots if we have less than 6 rows
    for j in range(plot_count, 6):
        fig.delaxes(axes[j])
        
    plt.tight_layout()
    plt.subplots_adjust(hspace=0.4)
    
    svg_out = "Figure_S2_Brain_Dosage_Grid.svg"
    png_out = "Figure_S2_Brain_Dosage_Grid.png"
    
    plt.savefig(svg_out, format="svg", bbox_inches="tight")
    plt.savefig(png_out, format="png", dpi=300, bbox_inches="tight")
    
    print(f"✅ Success! Generated 2x3 dosage grid for {plot_count} regions.")
    print(f"📁 Vector SVG: {svg_out}")
    print(f"📁 High-Res PNG: {png_out}")

if __name__ == "__main__":
    generate_figure_s2()
```
