## Supplemental Dataset 5 for "State-Dependent Transcriptomic Collapse of the Brain’s Lactate and Ketone Thermodynamic Sensors in Schizophrenia": README.pdf

**Supplemental Dataset 5: Subcortical Fine-Mapping and Disease Directional Validation**

**Study:** State-Dependent Transcriptomic Collapse of the Brain's Lactate and Ketone Thermodynamic Sensors in Schizophrenia

**Author:** Bryan A. Krantz

**Overview**

This repository contains the data, manual curation records, extraction logs, and processing scripts utilized to validate the directional transcriptomic collapse of the *HCAR* tandem array across subcortical and peripheral neural tissues (e.g., Nucleus Accumbens, Cerebellar Hemisphere, Spinal Cord).

While GTEx provides non-directional *P*-values, our peripheral pleiotropy analysis (Dataset 4) demonstrated that specific regulatory elements can invert their effect depending on the tissue environment. To rule out intra-brain enhancer pleiotropy, this dataset provides the manual directional validation confirming that the SCZ mutational burden strictly *downregulates* the metabolic governors across all evaluated neural circuits. This dataset generates the 5-panel dosage grid validating these thermodynamic failures (**Supplemental Figure S2**).

**Methodology: Manual Curation and Directional Alignment**

Standard Genotype-Tissue Expression (GTEx V8) data portals and APIs frequently obscure nominal eQTL data and Inverse Normal Transformed (INT) expression medians if they do not meet highly conservative, genome-wide FDR thresholds. To verify the biological direction of transcription, targeted manual curation was required.

1. **Manual Extraction:** Index SCZ risk variants were queried in the GTEx portal for subcortical and regional brain targets. Raw INT median expression values, exact genotype strings (e.g., CC, CT, TT), and sample sizes (n) were manually extracted from the UI violin plots via direct hover-state inspection to bypass automated masking.
2. **Disease Direction Alignment:** GTEx reports expression shifts strictly in terms of Reference vs. Alternate alleles. Custom Python scripts were utilized to cross-reference the extracted GTEx genotypes against the PGC Wave 3 Schizophrenia risk alleles. This logic dynamically oriented the raw expression deltas to the true pathogenic allele, calculating the "True Disease Delta."

**File Manifest**

**1. GWAS Query Targets (.tsv)**

These files contain the top index SNPs defining the 3' and 5' mutational skyscrapers, extracted directly from the PGC3 summary statistics. They serve as the reference for assigning the SCZ risk allele during directional calculations.

- HCAR_3prime_cluster_top_SNPs.tsv: Top variants residing within the *HCAR1* (lactate brake) regulatory domain.
- HCAR_5prime_cluster_top_SNPs.tsv: Top variants residing within the *HCAR2* (BHB cooling switch) regulatory domain.

**2. Raw Scrape Logs (.txt)**

- HCAR1_ENSG00000196917_6_rs28755851_chr12_123001735_A_T_b38_spinal_cord_scrape.txt: Raw GTEx extraction text utilized to validate the *HCAR1* spinal cord expression failure.
- HCAR2_ENSG00000182782_8_and_rs10734918_chr12_122436965_A_C_b38_other_brain_areas_scrape.txt: Raw GTEx extraction text utilized to validate the *HCAR2* subcortical failures (eQTLs in the Nucleus Accumbens, Hypothalamus, Cerebellum, etc.).

**3. Curated Transcriptomic Data (.xlsx)**

- cortex&other_brain_areas_GTEx_transcription_fine_data_medians_n_genotypes.xlsx: The master manual curation matrix. The other_brain_areas tab contains the hand-extracted INT medians, genotypes, and cohort sizes for the subcortical spot-checks.
- Subcortical_Disease_Direction_Calculated.xlsx: The finalized, computed dataset generated by the calculator script. This file merges the GWAS risk allele data with the GTEx expression data, providing the calculated directional shifts for plotting.

**4. Processing and Plotting Scripts (Python)**

- calculate_subcortical_disease_direction.py: The computational logic script that imports the manual curation matrix, dynamically identifies the SCZ risk allele, and calculates the true positive/negative transcriptomic shift to prove downregulation.
- figure_s2_subcortical_dosage.py: The visualization script utilizing matplotlib to generate the 2x3 dosage grids, graphically demonstrating the step-wise transcriptomic collapse of the metabolic receptors under increasing SCZ variant dosage.

**5. Output Figures**

- Figure_S2_Brain_Dosage_Grid.svg: Vector graphic output.
- Figure_S2_Brain_Dosage_Grid.png: High-resolution raster output.
